# SVEP1 couples adipose niche remodeling to macrophage activation and impaired metabolism in obesity

**DOI:** 10.64898/2026.09.19.751140

**Authors:** Nadav Kislev, Nahum Kavin, Roza Izgilov, Dafna Benayahu

## Abstract

The extracellular matrix (ECM) of adipose tissue regulates metabolic homeostasis, yet how ECM composition drives adipose dysfunction in obesity remains only partially defined. Multi-omics profiling of visceral adipose tissue identified the multidomain ECM protein SVEP1 as a candidate regulator of adipose remodeling; SVEP1 was elevated in obese human adipose tissue and correlated with insulin resistance and systemic inflammation, and interaction analysis predicted associations with ECM proteins, integrins, and cytoskeletal regulators. On a chow diet, *Svep1* heterozygous (*Svep1+/–*) mice gained less weight with lower fasting glucose, and their adipocytes showed enhanced lipolysis and AMPK activation. On a high-fat diet, *Svep1+/–* mice exhibited attenuated adiposity, improved glucose tolerance and insulin sensitivity, and fewer adipose macrophages with a shift away from a pro-inflammatory profile, alongside downregulation of inflammatory, lipid-metabolic, and ECM-remodeling pathways. Decellularized *Svep1*+/– matrix altered macrophage adhesion and phenotype relative to *Svep1+/+* matrix, identifying SVEP1 as a matrix-borne mediator coupling adipose remodeling to immune activation and metabolic impairment. Together, these findings position the adipose ECM as an active driver of immunometabolic dysfunction and nominate SVEP1 and its receptor network as candidate targets in obesity.

## Introduction

Adipose tissue is a metabolically active organ that continuously adapts to nutritional state, energy demand, and systemic inflammatory cues (*1*, *2*). During obesity, sustained caloric excess drives adipocyte hypertrophy, impairs lipid turnover, and causes progressive alterations in tissue architecture. The structural changes are tightly coupled to shifts in ECM composition, vascularization, and immune-cell infiltration, creating a microenvironment in which metabolic stress and inflammation reinforce one another. As adipocytes enlarge, local hypoxia, mechanical strain, and ECM remodeling promote the recruitment and activation of immune cells, particularly macrophages, which aggregate around dysfunctional adipocytes and modulate the tissue’s inflammatory and metabolic profiles (*3*, *4*). Together, these coordinated processes establish an adipose niche in which ECM-cell interactions, immune-stromal communication, and altered adipocyte function converge to shape systemic metabolic outcomes.

The ECM plays a central role in coordinating adipose tissue responses to metabolic stress. Beyond providing structural support, the ECM regulates adipocyte differentiation, lipid storage, and cellular viability. It also shapes the behavior of resident and infiltrating immune cells through biochemical and mechanical cues (*5*). Obesity induces marked alterations in ECM proteins, characterized by increased deposition of collagens and basement membrane components, as well as changes in cross-linking and tissue stiffness (*6*, *7*). These modifications, which culminate in fibrotic remodeling of the tissue, impair adipocyte expandability and restrict nutrient exchange, thereby reinforcing local inflammation (*8*, *9*). As a result, ECM remodeling does not merely accompany metabolic overload but actively determines adipose tissue homeostasis by orchestrating the interplay between stromal, adipocyte, and immune cells.

As matrix-dependent cues increase under metabolic stress, they broadly affect the immune cells in the tissue. The ECM alterations that constrain adipocyte expandability also regulate the trafficking, retention, and activation of immune cells through integrin-and other adhesion-dependent mechanisms (*10*, *11*). Modulating ECM-immune communication can reshape adipose tissue inflammation and thereby alter metabolic outcomes, establishing this axis as a major focus for understanding how structural remodeling translates into immunometabolic dysfunction (*12*, *13*). Within this framework, identifying specific ECM proteins that couple adhesive cues to immune cell behavior is essential for revealing new mechanistic regulators of adipose tissue.

SVEP1 (sushi, von Willebrand factor type A, EGF, and pentraxin domain-containing protein 1) has emerged as a multi-domain adhesion-associated ECM protein with properties that position it at the intersection of structural remodeling and inflammatory signaling. Originally characterized as a stromal cell-and basement membrane-associated adhesion molecule (*14*, *15*), SVEP1 is a known ligand of integrin α9β1 (*16*), integrin α4β1 (*17*), the receptor tyrosine kinase PEAR1 (*18*), and components of the ANG-TIE pathway, including Tie1 (*19*). Through these interactions, SVEP1 supports cell-matrix adhesion and influences migration, proliferation, and downstream signaling of immune and vascular cells (*20*). SVEP1 has also been implicated in cardiometabolic disease. Human genetic studies link *SVEP1* coding variants and elevated circulating protein levels to coronary artery disease, hypertension, and type 2 diabetes, and experimental models implicate SVEP1 in atherosclerotic plaque development and vascular inflammation (*21*, *22*). Although SVEP1 has been implicated in diverse inflammatory settings, its role in adipose tissue remains uncharacterized. Previous omics analyses indicate that SVEP1 is detectable in both visceral and subcutaneous adipose depots, and its expression is linked to metabolic and inflammatory traits (*23*, *24*). In immune contexts, SVEP1 influences monocyte differentiation and macrophage behavior via integrin-dependent signaling, suggesting potential relevance to adipose tissue macrophages (*20*, *25*). Despite these observations, the contribution of SVEP1 to ECM-immunometabolic interactions and its role in obesity-induced adipose tissue remodeling have not been investigated.

In this study, we combined integrated multi-omics profiling with in vivo mouse genetic models and functional cellular assays to delineate the role of SVEP1 in adipose tissue regulation. Using parallel transcriptomic and proteomic analysis of visceral adipose tissue, we identified SVEP1 as an ECM component upregulated under a high-fat diet (HFD). We characterized *SVEP1* expression across adipose compartments in mice and human cohorts and constructed the SVEP1 protein-protein interaction network and receptor landscape. Using *Svep1*+/– mice, we showed that reduced *SVEP1* expression attenuates HFD-induced adiposity, improves glucose tolerance, enhances adipocyte metabolic capacity, and reprograms adipose macrophages away from a pro-inflammatory state. Bulk RNA-seq of *Svep1*+/– adipose tissue, combined with pathway modeling and receptor-pathway intersection analysis, revealed coordinated suppression of integrin-linked and immune signaling programs. Using adipocyte-macrophage co-cultures and decellularized ECM, we found that matrix produced by *Svep1*-deficient cells is altered in its capacity to support macrophage adhesion, morphology, and inflammatory gene expression. Together, these findings define SVEP1 as an ECM regulator that shapes tissue remodeling and engages immune-cell receptors, with consequences for metabolic function during obesity.

## Methods and Materials

### Animal models, diet-induced obesity, and tissue collection

*Svep1* heterozygous knockout mice (*Svep1*+/–) were originally generated through the Knockout Mouse Project (KOMP) (kindly provided by the Stitziel laboratory, Washington University in St. Louis, USA) (*18*, *20*). Male C57BL/6J mice (6 weeks of age) were either fed a standard chow diet (CHD) or a 60% HFD (Research Diet, D12492) for 12 weeks. Mice were housed under standard conditions with a 12 h light/dark cycle, controlled temperature (22 ± 1°C), and free access to food and water. At the indicated time points, mice were fasted, anesthetized, and epididymal and inguinal adipose depots were collected, either processed fresh or snap-frozen in liquid nitrogen and stored at −80°C. All procedures were approved by the Tel Aviv University Institutional Animal Care and Use Committee (IACUC) and performed in accordance with institutional guidelines (protocol number TAU-MD-IL-2212-179-4).

### Glucose and insulin tolerance tests

For glucose tolerance tests (GTT), mice were fasted for 12–18 h with free access to water and then injected intraperitoneally with D-glucose (2 g/kg body weight; Sigma-Aldrich, G8270). For insulin tolerance tests (ITT), mice were fasted for 6 h and injected intraperitoneally with human insulin (0.75 U/kg; MP Biomedicals, SKU:02193900-CF). Blood glucose was measured from tail vein samples at 0, 15, 30, 60, and 120 min using a handheld glucometer (Contour^®^ Plus; Bayer).

### Stromal vascular fraction isolation

The stromal vascular fraction (SVF) was isolated from inguinal and epididymal adipose depots. Tissues were minced and transferred into cell isolation buffer consisting of Hanks’ balanced salt solution (HBSS) supplemented with 2% fetal bovine serum (FBS). Samples were digested at 37°C for 20–30 min with collagenase type I (1 mg/ml; Worthington Biochemical, LS004196) and dispase (Sigma, 165859) in isolation buffer under gentle agitation. The digested material was diluted with an equal volume of isolation buffer, filtered through sterile gauze, and centrifuged at 400×g for 5 min at room temperature. Floating mature adipocytes were removed, and the SVF pellet was washed twice with isolation buffer. Red blood cells were lysed using RBC lysis buffer (Sartorius 01-888-1B) for 3 min, then the suspension was diluted and centrifuged (2,500 rpm, 5 min). The final pellet was passed through a 40-µm cell strainer and resuspended in the appropriate buffer or medium for downstream analyses.

### Adipose-derived mesenchymal stromal cells and adipogenic differentiation

Isolated SVF cells from murine inguinal adipose tissue were plated in growth medium (DMEM, 4.5 g/L glucose; Gibco, 11965) supplemented with 20% FBS (HyClone SH30071.03), 1% L-glutamine (Biological Industries 03-020-1A), 0.1% penicillin-streptomycin (Sigma-Aldrich P3032/85555), and 0.5% HEPES (Biological Industries 03-025-1B), and allowed to adhere; non-adherent cells were removed after 24 h. Adipose-derived mesenchymal stromal/progenitor cells (AMSCs) were expanded until confluence. For adipogenic differentiation, confluent cultures were induced with differentiation medium containing insulin (10 µg/ml), dexamethasone (1 µM), IBMX (400 µM), and rosiglitazone (1 µM) in DMEM with 10% FBS for 48 h, followed by maintenance medium with insulin alone for 4–10 days, with medium changes every 2–3 days. The level of adipogenesis (LOA) was quantified from stitched macroscopic images (*26*).

### Bone marrow isolation and macrophage differentiation

Bone marrow cells were harvested from femurs and tibias of wild-type C57BL/6J mice. After removal of attached muscle, bones were cut at the joints and flushed with ice-cold phosphate-buffered saline (PBS) into a 50-ml tube. The cell suspension was filtered through a 40-µm strainer and centrifuged at 200×g for 5 min. For bone marrow-derived macrophage (BMDM) cultures, cells were plated in growth medium (DMEM, 4.5 g/L glucose) supplemented with 10% FBS, 1% L-glutamine, 0.1% penicillin-streptomycin, and 0.5% HEPES. Differentiation into macrophages was induced by adding 10% CMG-conditioned medium as a source of M-CSF, with medium changes twice weekly for 7 days; macrophage identity was confirmed by morphology and F4/80 expression.

### Adipocyte-macrophage co-culture

For co-culture experiments, AMSCs from *Svep1*+/+ or *Svep1*+/– mice were differentiated into adipocytes as described above. On day 14, differentiated adipocyte cultures were washed, and BMDMs were plated on the adipocyte layer in DMEM with 10% FBS. Co-cultures were stimulated with LPS (100 ng/ml; Sigma-Aldrich, L4516) for 8 h. At the end of the incubation period, cells were harvested for RNA isolation and qPCR analysis of adipogenic, metabolic, and inflammatory gene expression.

### Generation of decellularized extracellular matrices (dECM)

Primary adipose-derived stromal cells from *Svep1*+/+ or *Svep1*+/– mice were cultured to confluence as undifferentiated cells to establish an SVEP1-containing ECM layer. Cultures were washed twice with PBS and subjected to three freeze-thaw cycles at −80°C, followed by incubation in decellularization buffer (PBS containing 0.5% Triton X-100 and 20 mM NH₄OH) for 30 min at room temperature. Residual DNA was removed by DNase I treatment (100 U/ml; Sigma-Aldrich DN25) for 30–60 min at 37°C. Matrices were washed extensively with PBS and either used immediately for downstream assays or stored short-term in PBS at 4°C.

### Macrophage adhesion and morphology assays on decellularized matrices

BMDMs were detached, counted, and seeded onto decellularized matrices derived from *Svep1*+/+ or *Svep1*+/– stromal cells in 96-well plates at a density of 5×10⁴ cells/well in serum-containing medium. After 30–60 min at 37°C, non-adherent cells were removed by gentle washing with PBS. Adherent cells were fixed in 4% paraformaldehyde for 15 min, stained with 4ʹ,6-diamidino-2-phenylindole (DAPI) and phalloidin, and imaged using **an**EVOS FL Auto 2 microscope (Thermo Fisher Scientific). The number of adherent cells per field and morphological parameters (cell area, perimeter, and circularity) were quantified in ImageJ/Fiji.

### Protein extraction and western blotting

Cells were lysed in RIPA buffer supplemented with protease inhibitors (Sigma-Aldrich P8340) and phosphatase inhibitors (Sigma-Aldrich P0044). Protein concentration was determined by bicinchoninic acid (BCA) assay (Pierce/Thermo 23225). Equal amounts of protein (30 µg per lane) were resolved by SDS-PAGE on 10% polyacrylamide gels and transferred to PVDF membranes. Membranes were blocked in 5% non-fat dry milk in TBS-T for 1 h at room temperature, then incubated overnight at 4°C with primary antibodies against: phospho-AKT (Ser473; Cell Signaling Technology, 4060, 1:1000); total AKT (Cell Signaling Technology, 4691, 1:1000); phospho-AMPKα (Thr172; Cell Signaling Technology, 2535, 1:1000); total AMPKα (Cell Signaling Technology, 5831, 1:1000); UCP1 (Abcam ab10983, 1:1000); and β-actin or GAPDH as loading control (Santa Cruz SC-47778 (β-actin clone C4), 1:1000). After incubation with HRP-conjugated secondary antibodies (Jackson 111-035-003 (anti-rabbit) and 115-035-003 (anti-mouse), 1:10,000), bands were visualized using ECL reagent (Pierce SuperSignal West Pico PLUS, 34580) on a Fusion FX7 (Vilber) imaging system. Band intensities were quantified using ImageJ.

### Lipolysis and glycerol assays

Lipolysis was induced in differentiated adipocytes by incubation in serum-free medium containing forskolin (10 µM; Thermo Fisher Scientific, J63292) for 2 h under fasting conditions. Glycerol release into the conditioned medium was quantified as a readout of lipolytic activity using a colorimetric glycerol assay kit (Promega Glycerol-Glo Assay, J3151). Values were normalized to total protein content measured by BCA assay.

### Immunofluorescence staining and confocal imaging

For whole-mount staining of adipose tissue, freshly isolated epididymal fat pads were fixed in 4% paraformaldehyde for 2 h at room temperature, washed with PBS, and blocked in PBS containing 0.3% Triton X-100 and 5% normal goat serum for 2 h at room temperature. Samples were incubated overnight at 4°C with primary antibodies: anti-SVEP1, lab-generated (*15*, *27–29*), 1:100; anti-Perilipin-1/PLIN1 (sc-390169, Santa Cruz Biotechnology, 1:100); anti-F4/80 (sc-377009, Santa Cruz Biotechnology, 1:100); anti-CD11b (03221-65, PE/Cy5, the same conjugated antibody used for flow cytometry; Biogems, 1:100); anti-ITGA9 (sc-71428, Santa Cruz Biotechnology, 1:100); anti-Nidogen-1/NID1 (MAB1946, clone ELM1; Sigma-Aldrich/Millipore, 1:100). After washing, tissues were incubated with the appropriate Alexa Fluor-conjugated secondary antibodies (A-21127, AF555 anti-mouse IgG1; and A-21141, AF488 anti-mouse IgG2b; Thermo Fisher Scientific, 1:500) for 2 h at room temperature, then mounted in DAPI-containing mounting medium (Fluoroshield with DAPI, EMS #17985-10). Images were acquired on a Leica SP8 confocal microscope. For lipid droplet visualization, adipose tissues or cultured cells were stained with Nile Red (1 µg/ml; Sigma-Aldrich, N3013) for 15 min, then imaged using appropriate filter sets. For cultured cell immunofluorescence (pFAK), cells were fixed, blocked, and stained as above using anti-phospho-FAK (Tyr397; sc-81493, Santa Cruz, clone 14; 1:100).

### Image analysis

All image-based quantifications were performed in ImageJ/Fiji. For adipocyte morphology in whole-mount and paraffin tissue sections, cell area was quantified using the Analyze Particles function on threshold fluorescence images. Lipid droplet size, number, and area per cell were quantified using the Trainable WEKA Segmentation plugin (*30*), with measurements normalized to cell number. SVEP1 fluorescence intensity along adipocyte membranes was measured in ImageJ, and CD11b+ cell and macrophage counts per field were determined by automated particle counting on threshold images. Corrected total cell fluorescence (CTCF) of pFAK and macrophage morphology on dECM (cell area, perimeter, circularity) were also quantified in ImageJ.

### Flow cytometry

SVF cells isolated from epididymal adipose tissue were resuspended in staining buffer (PBS with 2% FBS) and incubated with 2% goat serum + 0.1% NaN3 for 10 min on ice. Cells were then stained with the following fluorophore-conjugated antibodies for 30 min at 4°C (antibodies listed in Table S2): anti-CD45 (clone 30-F11; BioLegend 103128 [AF700]), anti-F4/80 (clone BM8.1; Biogems 02922-80 [FITC]), anti-CD11b (clone M1/70; Biogems 03221-65 [PE/Cy5]), anti-MHCII (clone M5/114.15.2; BioLegend 107629 [PE/Cy7]), anti-CD11c (clone N418; BioLegend 117323 [APC/Cy7]), anti-ITGA9 (polyclonal goat IgG, immunogen Tyr31-Val979; R&D Systems FAB3827P [PE]). After washing, cells were resuspended in staining buffer containing DAPI for viability exclusion and acquired on a CytoFLEX 5L (Beckman Coulter) flow cytometer. Data were analyzed using Kaluza software (Beckman Coulter). Adipose tissue macrophages (ATMs) were gated as CD45+CD11b+F4/80+ cells, and subpopulations were defined by major histocompatibility complex class II (MHCII) and CD11c expression (*31*). ITGA9 mean fluorescence intensity (MFI) was recorded within each ATM subset.

### RNA isolation and quantitative PCR (qPCR)

Total RNA was isolated from tissues and cultured cells using the Hybrid-R RNA isolation kit (GeneAll, 305-101). RNA concentration and quality were assessed by NanoDrop. Complementary DNA (cDNA) was synthesized using the Ultrascript cDNA synthesis kit (PCR Biosystems, PB30.21-10) with random hexamer primers. Quantitative PCR was performed using SYBR Green-based reagents (qPCR BIO SyGreen Mix Lo-ROX; PCR Biosystems, PB20.11-05) on a StepOne Plus Real-Time PCR System (Thermo Fisher Scientific). Gene expression was normalized to *Hprt* for macrophages and to *Rplp0* for adipocytes and adipose tissue samples using the ΔΔCt method. Primer sequences are listed in Table S1.

### Bulk RNA sequencing and transcriptomic analysis

Total RNA from epididymal adipose tissue was isolated with the Hybrid-R kit, and RNA integrity was confirmed on a TapeStation 4200 instrument (Agilent) before library construction. Sequencing libraries were built using a 3ʹ-tag protocol incorporating unique molecular identifiers (UMIs). Adapter and poly-A/T sequences were removed with cutadapt (*32*), fragments below 30 bp were excluded, and the surviving reads were aligned to 1,000-base 3ʹ-UTR windows of the mouse reference genome (GRCm38/mm10, RefSeq) using STAR in EndToEnd mode with outFilterMismatchNoverLmax set to 0.05 (*33*). Duplicate reads — those sharing both a gene assignment and a UMI — were collapsed, and per-gene counts were then tabulated with htseq-count (*34*). Differential expression was evaluated in DESeq2 using the Wald test, with betaPrior disabled; gene-wise p-values were adjusted using Independent Hypothesis Weighting (IHW; FDR < 0.05), and transcripts were additionally required to exceed an effect-size cut-off of |log₂FC| ≥ 0.5 to be considered differentially expressed genes (DEGs), with the same criteria applied to both RNA-seq contrasts (*35*).

Pathway enrichment analysis of DEGs was performed using clusterProfiler for Gene Ontology Biological Process (GOBP), Kyoto Encyclopedia of Genes and Genomes (KEGG), and Reactome terms (FDR < 0.05) (*36*). Enrichment results were visualized using enrichment maps and ClueGO (*37*) in Cytoscape v3.10 (*38*), with terms grouped based on shared gene membership. Transcription factor (TF) activity was inferred using DoRothEA (confidence levels A–C) and decoupleR, applying the multivariate linear model (MLM) method (*39*). Pathway activity was modeled with PROGENy (*40*). Ligand-receptor signaling programs were analyzed using BulkSignalR, restricting to curated ligand-receptor pairs and summarizing pathway-level regulation across conditions (*41*). Macrophage polarization state was evaluated using MacSpectrum, which provides a Macrophage Polarization Index (MPI) and Activation-Induced Macrophage Differentiation Index (AMDI) (*42*). Cell-type composition was estimated by deconvolving bulk RNA-seq data using single-nucleus reference profiles from the adipose atlas (GSE176171) via a regression-based deconvolution framework (*43*). Multi-source pathway analysis integrating WikiPathways, GeneGo (MetaCore), Reactome, PubChem, QIAGEN, R&D Systems, and Cell Signaling Technology was performed using GeneAnalytics (*44*).

### LC-MS/MS proteomics

For proteomic profiling, epididymal adipose tissue from CHD-and HFD-fed mice was processed for liquid chromatography-tandem mass spectrometry (LC-MS/MS) as previously described (*45*). Briefly, tissues were homogenized in urea-based lysis buffer, protein concentration was determined by BCA assay, and 50 µg per sample was reduced, alkylated, and digested overnight with sequencing-grade trypsin. Peptides were desalted, separated by reverse-phase nano-LC, and analyzed on a Q Exactive HF Orbitrap mass spectrometer (Thermo Fisher Scientific). Raw data were processed with MaxQuant against the UniProt Mus musculus reference proteome, and differentially abundant proteins (DAPs) were identified using moderated t-tests in Perseus with Benjamini–Hochberg correction (FDR < 0.05). Functional enrichment of DAPs was performed using Reactome pathway analysis and GSEA (fgsea package in R), and GOCC summed abundances were calculated by aggregating LFQ intensities of proteins annotated to each compartment (*46*).

### SVEP1 protein-protein interaction network and receptor mapping

The human SVEP1 protein structure was obtained from the AlphaFold Protein Structure Database (UniProt ID: Q4LDE5) (*47*). The SVEP1 protein-protein interaction (PPI) network was constructed using the PrePPI database (*48*), which combines structural modeling with functional, evolutionary, and expression-based evidence. Interactors with probability scores above the recommended confidence threshold were retained as the SVEP1 interactome. Gene Ontology Cellular Component (GOCC) and pathway enrichment analyses (Reactome, KEGG) were performed in R using the clusterProfiler and ReactomePA packages (FDR < 0.05) (*36*, *49*). Enrichment results were visualized and grouped into functionally related clusters using ClueGO (*37*) in Cytoscape. UniProtKB annotations for receptors and membrane-associated proteins were used to identify SVEP1-associated receptors within this interactome (28 candidate receptors identified by cross-referencing with the UniProt human receptome, n = 1,656 curated receptor proteins). Multi-source pathway scores for each receptor were obtained using GeneAnalytics across seven pathway resources (WikiPathways, MetaCore, Reactome, QIAGEN, CST, R&D Systems, and PubChem) (*44*). Receptor-pathway associations were integrated with DEG-derived pathway results from bulk RNA-seq analyses to generate intersection matrices, and receptor-centered signaling networks were visualized in Cytoscape (*38*).

### Human transcriptomic datasets and meta-analysis

Bulk tissue *SVEP1* expression across human tissues was assessed using the GTEx Portal (GTEx Analysis Release V11; dbGaP accession phs000424.v11.p2) (*50*). Expression values were retrieved as normalized TPM values, log2-transformed for visualization, and summarized across tissues in R. For cell-type-resolved analyses, normalized and annotated human and mouse adipose single-nucleus RNA-seq data were obtained from the Emont et al. adipose atlas (*43*) (GSE176171). *SVEP1* expression was summarized across annotated adipocytes, adipose stem and progenitor cells (ASPCs), endothelial cells, lymphatic endothelial cells (LECs), macrophages, monocytes, T cells, and B cells, and displayed according to the stratifications used in the corresponding atlas analyses. Cross-cohort validation of *SVEP1* expression in obese versus non-obese human adipose tissue was performed using the meta-analysis functionality of www.adiposetissue.org (*51*). Correlations between *SVEP1* expression and anthropometric or metabolic traits were obtained from the clinical module portal of www.adiposetissue.org (*51–73*). The portal integrates thousands of human adipose transcriptomic cohorts and reports pooled standardized mean differences (SMDs) with 95% confidence intervals under both common-effect and random-effects models, together with heterogeneity statistics including I² and τ².

### Schematic illustrations

Schematic illustrations were created in BioRender (https://BioRender.com).

### Statistical analysis

Statistical analyses were performed in GraphPad Prism (version 9.1.1) and R (version 4.2 or later). Data are presented as mean ± standard error of the mean (SEM) unless otherwise specified. Normality was assessed using the Kolmogorov–Smirnov test, and outliers were identified using the ROUT method (Q = 1%). Comparisons between two groups were made using two-tailed unpaired Student’s t-tests or Mann–Whitney tests for non-normally distributed data. For comparisons among more than two groups, one-way or two-way analysis of variance (ANOVA) was used with Tukey or Sidak post hoc tests as appropriate, or Kruskal–Wallis with Dunn’s post-test for non-parametric data. Glucose and insulin tolerance test curves were analyzed by two-way repeated-measures ANOVA with Sidak correction; the corresponding area under the curve (AUC, glucose tolerance tests) and area over the curve (AOC, insulin tolerance tests) values were compared between genotypes by two-tailed unpaired Student’s t-test or Mann–Whitney test, as appropriate. For omics-level analyses, multiple-testing correction was performed using the Benjamini–Hochberg method, and a false discovery rate (FDR) < 0.05 was used as the significance threshold. For all other analyses, a p-value < 0.05 was considered statistically significant.

## Results

### Integrated multi-omics analysis identifies extracellular matrix remodeling as a hallmark of adipose tissue dysfunction in diet-induced obesity

To define the molecular programs shaping visceral adipose tissue responses to nutritional challenge, six-week-old male mice were maintained on either a CHD or a HFD for 12 weeks. Epididymal adipose tissues were then subjected to parallel bulk RNA-seq and LC-MS/MS proteomics, followed by integrated pathway analysis, transcription factor activity profiling, and ligand-receptor analyses to identify the dominant regulatory signals induced by HFD (Fig. 1a). Differential expression analysis demonstrated a robust transcriptional response between diets (Fig. S1a), highlighting 1,736 significantly regulated genes (770 upregulated and 966 downregulated under HFD) and representing the broad scope of adipose tissue remodeling induced by nutritional excess. Reactome pathway enrichment of the transcriptomic data (Fig. 1b) confirmed prominent activation of ECM organization, cytoskeletal remodeling, and immune modulation, while highlighting marked suppression of respiratory electron transport and amino acid metabolism. GOBP analysis (Fig. 1c) further revealed coordinated induction of leukocyte activation, cytokine signaling, and extracellular structural organization alongside consistent downregulation of oxidative phosphorylation and ATP-generating programs, with heatmaps of representative genes illustrating elevated expression of collagens, basement membrane components, chemokines, and immune receptors contrasted with reduced mitochondrial and metabolic transcripts.

**Figure 1:**
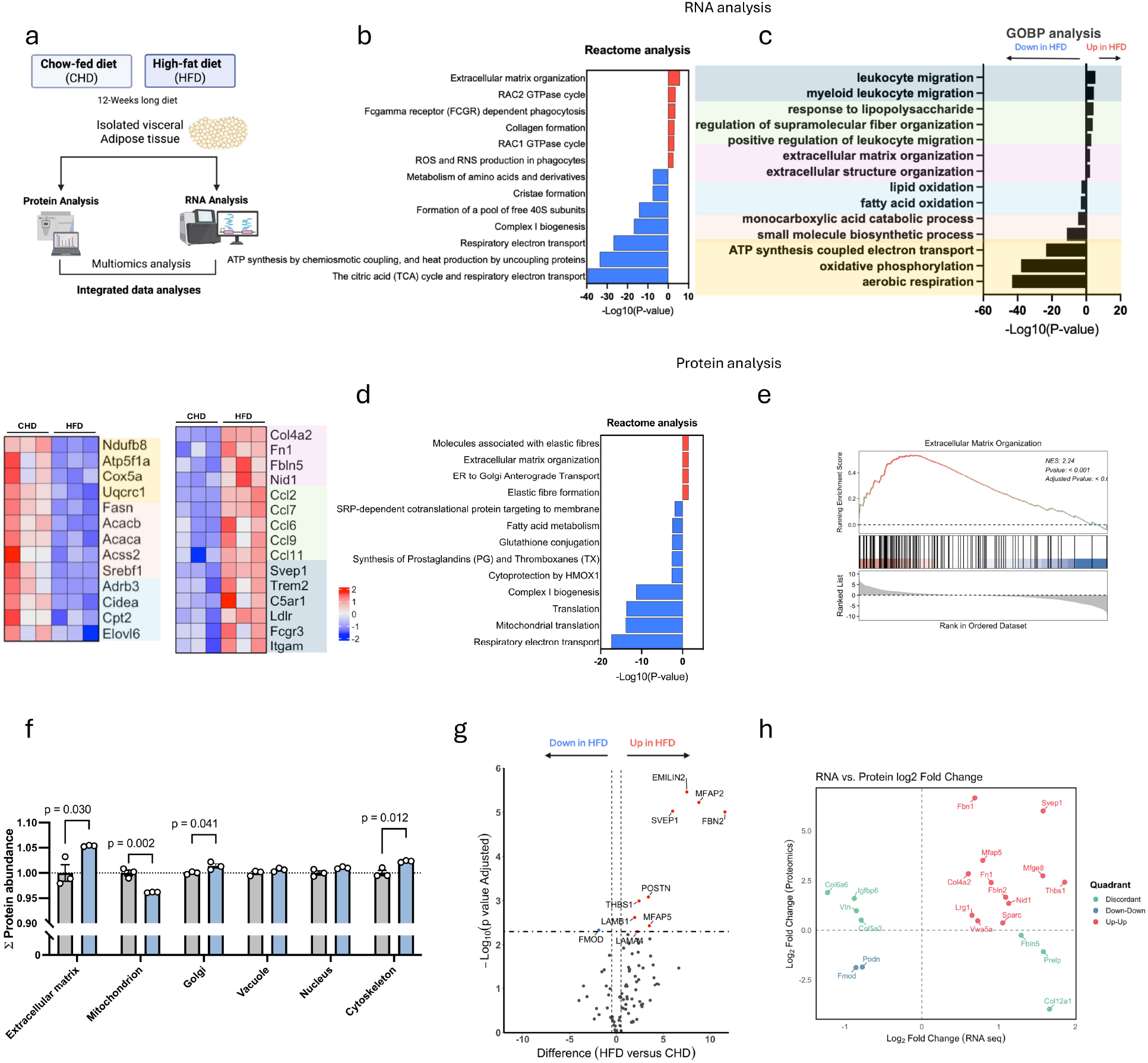
Integrated multi-omics analysis identifies extracellular matrix remodeling in visceral adipose tissue under high-fat diet. (a) Schematic overview of the experimental and analytical pipeline. Epididymal visceral adipose tissue was collected from CHD-fed and HFD-fed mice (12 weeks; n = 3 per group) andsubjected to parallel RNA-seq and LC-MS/MS proteomic analyses followed by integrated bioinformatic analyses. (b) Reactome pathway enrichment analysis of DEGs. (c) GOBP enrichment analysis of DEGs, with representative heatmaps of genes contributing to selected downregulated (left) and upregulated (right) pathways (n = 3 per group). (d) Reactome pathway enrichment analysis of DAPs (FDR < 0.05). (e) Gene set enrichment analysis (GSEA) of the Extracellular Matrix Organization pathway in HFD samples (NES and FDR as indicated). (f) Relative protein abundance across selected GOCC categories, including extracellular matrix, mitochondrial organization, Golgi, vacuole, nucleus, and cytoskeleton, in CHD (gray) and HFD (blue) visceral adipose tissue (n = 3 per group). Statistical significance was assessed using a two-tailed unpaired Student’s t-test. (g) Volcano plot of ECM-associated proteins highlighting significantly altered components in HFD versus CHD. (h) Correlation analysis of matched ECM-associated transcripts and proteins (RNA vs. protein log₂ fold change), indicating concordantly upregulated (red), concordantly downregulated (blue), and discordant (green) molecules. Data are presented as mean ± SEM.

To investigate upstream regulatory drivers, we first applied PROGENy (Fig. S1b), a footprint-based method that infers pathway activity from the expression of well-characterized downstream target genes. This analysis revealed activation of NF-κB, PI3K, MAPK, hypoxia, and TGF-β programs together with attenuation of EGFR and androgen signaling in HFD adipose tissue. Volcano plots of NF-κB-and EGFR-responsive genes (Fig. S1c) further supported these opposing pathway shifts, demonstrating induction of canonical inflammatory targets (e.g., *Ccl2*, *Tnfaip3*, *Ptgs2*) alongside repression of EGFR-associated genes (e.g., *Egr1*, *Phlda1*, *Dusp5*). DoRothEA, which infers transcription factor activity from curated sets of target genes (regulons) (Fig. S1d), identified SPI1, CEBPB, and NF-κB-related factors as the dominant activated regulators, while metabolic regulators such as SREBF1/2 and PPAR-related factors were suppressed. Target gene analysis (Fig. S1e) revealed robust induction of SPI1-driven inflammatory genes and coordinated repression of SREBF1-dependent lipogenic programs, reinforcing the shift from metabolic to immune transcriptional control. BulkSignalR, which infers ligand-receptor interactions and their downstream pathway support from bulk expression data (Fig. S1f), demonstrated enrichment of integrin signaling, chemokine-mediated communication, and ECM-immune crosstalk under HFD conditions.

We next assessed how these transcriptional shifts were reflected at the protein level. Reactome analysis of differentially abundant proteins (Fig. 1d, S2a) closely mirrored the transcriptomic results, showing strong upregulation of ECM organization, ER-to-Golgi transport, and immune-associated modules, as well as a marked reduction in mitochondrial translation and oxidative phosphorylation. Gene set enrichment analysis confirmed a coordinated induction of the Extracellular Matrix Organization pathway in HFD samples (Fig. 1e, S2b). Quantification of relative protein abundance across GOCC categories (Fig. 1f) demonstrated significant increases in ECM-associated proteins, vesicular and cytoskeletal fractions, and decreases in mitochondrial proteins. Within the ECM, several proteins were significantly upregulated under HFD, including SVEP1, EMILIN2, MFAP2, and FBN2, reflecting coordinated changes in structural and matricellular components (Fig. 1g). To determine whether these protein-level changes were mirrored at the transcript level, the RNA-seq and proteomic datasets were integrated. Correlation analyses of matched RNA-protein pairs (Fig. S2c) and enrichment of concordantly regulated pairs (Fig. S2d) identified a dominant transcriptional-proteomic signature of HFD-induced remodeling, characterized by activation of cell adhesion and ECM interaction pathways and suppression of oxidative and mitochondrial processes. Focusing on matched ECM gene-protein pairs (Fig. 1h) further delineated the ECM compartment into concordantly upregulated, downregulated, and discordant molecules. Notably, SVEP1 emerged as a concordantly upregulated ECM component at both transcriptional and proteomic levels, suggesting its potential role as a mediator of obesity-driven adipose ECM remodeling.

### SVEP1 is enriched in adipose stromal compartments and associates with metabolic dysfunction

Following the identification of SVEP1 as a leading candidate in the multi-omics screen, its expression in adipose tissues was further characterized. Quantitative PCR analysis of *Svep1* expression in the SVF and in isolated adipocytes from epididymal (visceral) adipose tissue of chow-fed mice (Fig. 2a) demonstrated enriched expression in adipocytes. *Svep1* mRNA levels were significantly higher in epididymal adipose tissue from HFD-fed mice compared with chow-fed controls (Fig. 2b), supporting the multi-omics findings. Whole-mount immunofluorescence staining of epididymal adipose tissue for SVEP1, perilipin, and DAPI (Fig. 2c) revealed the spatial localization of SVEP1 at the adipocyte surface and extracellular matrix, and showed increased staining intensity in HFD-fed mice relative to chow-fed mice.

**Figure 2.**
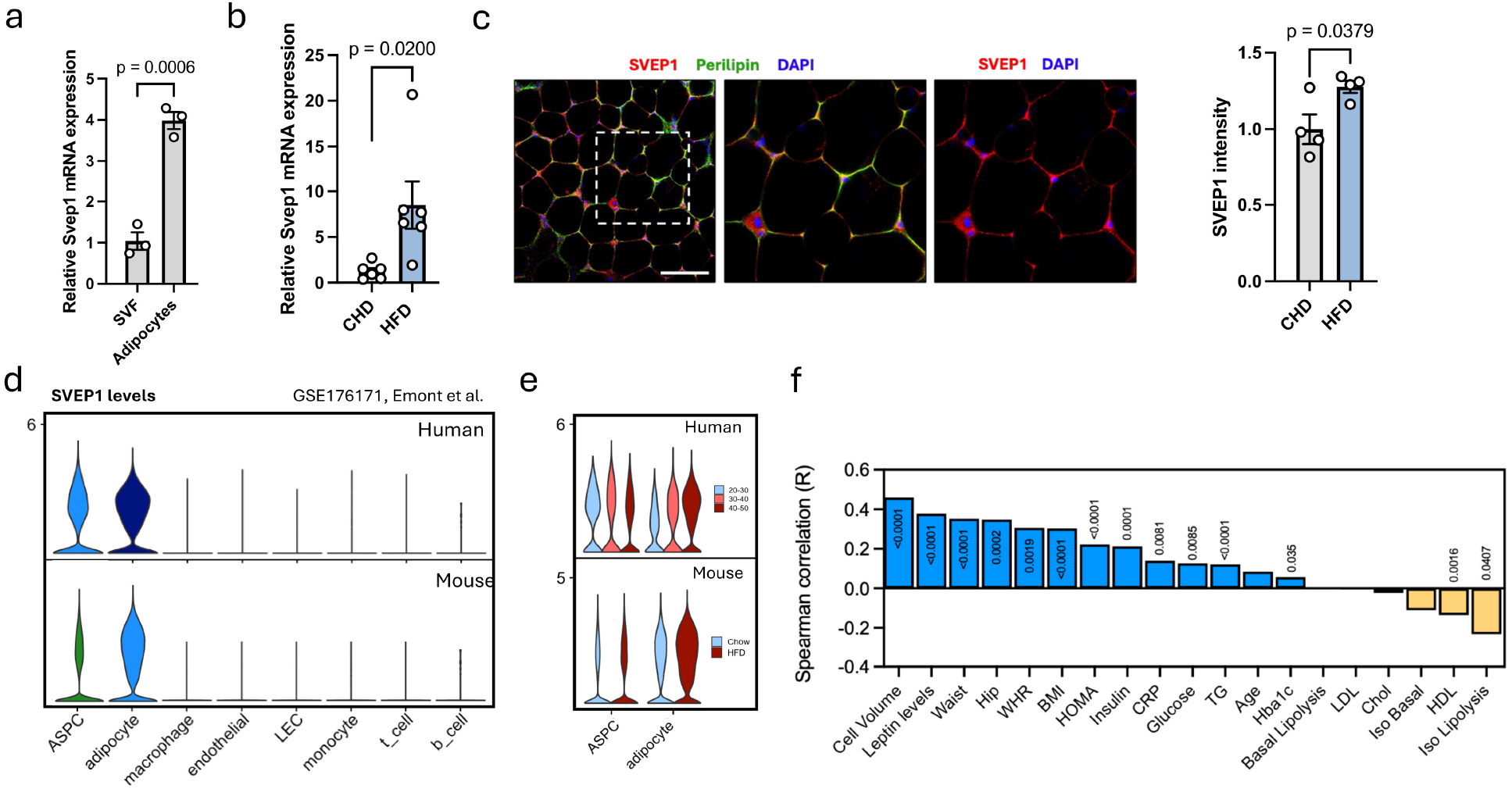
SVEP1 is enriched in adipose stromal compartments and associates with metabolic dysfunction. (a) qPCR analysis of *Svep1* mRNA expression in SVF and primary adipocytes isolated from visceral adipose tissue of chow-fed mice (n = 3). (b) qPCR analysis of *Svep1* mRNA expression in epididymal adipose tissue from CHD (gray)-and HFD (blue)-fed mice (n = 6 per group). (c) Representative whole-mount immunofluorescence images of visceral adipose tissue stained for SVEP1 (red), perilipin (green), and DAPI (blue), with quantification of SVEP1 fluorescence intensity (n = 4 mice). Scale bars = 166 µm. (d) *SVEP1* expression across human (top) and mouse (bottom) adipose stromal and immune cell populations from the Emont et al. adipose single-cell RNA-seq dataset (GSE176171), including ASPCs, adipocytes, macrophages, endothelial cells, LECs, monocytes, T cells, and B cells. (e) *SVEP1* expression across human adipose stromal and immune cell populations using the Emont et al. single-cell RNA-seq, stratified by BMI group (20–30, 30–40, and 40–50; top) and, in mice, by diet condition (chow versus HFD; bottom). Cell types include ASPCs and adipocytes. (f) Spearman correlation analysis of *SVEP1* expression with anthropometric and metabolic traits, including adipocyte cell volume, leptin, waist circumference, hip circumference, waist-to-hip ratio, BMI, HOMA-IR, insulin, CRP, glucose, triglycerides, HbA1c, basal lipolysis, isoproterenol-stimulated lipolysis, and circulating lipid measures. Correlation coefficients (R) are shown, with p values indicated above bars. Statistical significance for comparisons between two groups was assessed using two-tailed unpaired Student’s t-test or Mann–Whitney test, as appropriate. Data are presented as mean ± SEM.

*SVEP1* expression was then evaluated across more than 30 human tissues using the GTEx dataset (Fig. S3a), revealing that *SVEP1* is most highly expressed in adipose tissue among tissues enriched in ECM and stromal content. At single-cell resolution, *SVEP1* expression was assessed using human and mouse adipose single-nucleus RNA sequencing (snRNA-seq) datasets (Fig. 2d), which demonstrated high expression levels in ASPCs. Stratification by body mass index (BMI) group and diet condition (Fig. 2e) indicated preferential upregulation of *SVEP1* in adipocytes from obese individuals and from HFD-fed mice. These findings support a cell-state-dependent induction of *SVEP1* under metabolic stress.

To assess human clinical relevance, a meta-analysis of *SVEP1* expression was performed across 20 studies comparing obese (n = 1,014) and non-obese (n = 1,505) subjects (Fig. S4). The meta-analysis module of adiposetissue.org, a curated knowledge portal that aggregates *SVEP1* expression data from public human adipose transcriptomic cohorts and computes standardized mean differences under common-and random-effects models (*51*), was employed. The pooled analysis demonstrated significant upregulation of *SVEP1* in obese adipose tissue under both common-effect and random-effects models. Spearman correlation analysis of *SVEP1* expression with anthropometric and metabolic parameters (Fig. 2f) identified significant positive associations with adipocyte volume, leptin, waist circumference, hip circumference, waist-to-hip ratio, BMI, homeostatic model assessment of insulin resistance (HOMA-IR), fasting insulin, and C-reactive protein (CRP), as well as inverse associations with high-density lipoprotein (HDL) cholesterol and with basal and isoproterenol-stimulated lipolysis. These results indicate that higher *SVEP1* expression is associated with reduced adrenergic lipolytic capacity. This convergent pattern, spanning markers of adiposity, insulin resistance, impaired lipid mobilization, and systemic inflammation, suggests that SVEP1 serves not only as a correlate of tissue expansion but also as a potential functional node linking ECM remodeling to adipose metabolic dysfunction.

### The predicted SVEP1 interactome links ECM organization with receptor-mediated immune signaling

Having established that SVEP1 is induced in adipose tissue and tracks with metabolic dysfunction, we next asked how it might act, as no receptor or mechanism has been defined for SVEP1 in this tissue. We therefore used its domain architecture and predicted interactome to generate mechanistic hypotheses and to identify candidate interactors and pathways. SVEP1 is a large multidomain protein that contains CCP/sushi repeats, EGF-like domains, a vWA domain, and a C-terminal pentraxin domain (Fig. 3a). This modular architecture indicates broad extracellular interaction potential and motivates a systematic investigation of its predicted interactome. To characterize the functional interaction landscape of SVEP1 in adipose tissue, a protein-protein interaction network was constructed using the PrePPI framework, which integrates structural, functional, evolutionary, and expression-based evidence. GOCC classification of over 120 predicted interactors demonstrated strong enrichment in the extracellular region, basement membrane, integrin complex, and secretory compartments (Fig. 3b). Functional pathway enrichment analysis revealed significant overrepresentation of ECM organization, integrin-mediated cell surface interactions, complement activation, collagen biosynthesis, and cytoskeletal regulation in both KEGG and Reactome frameworks (Fig. 3c). ClueGO clustering (Fig. 3d) grouped SVEP1-associated proteins into distinct functional modules, including the ECM, cell-substrate junctions, integrin complexes, lysosomal/secretory compartments, and membrane microdomains. To determine the cell-type distribution of the SVEP1 interactome in adipose tissue, expression of the 122 predicted interactors was examined in a human adipose snRNA-seq dataset (Fig. S5a, (*43*)). The interactome segregated into major stromal/ASPC ECM, endothelial/lymphatic endothelial, and macrophage/monocyte immune clusters, positioning SVEP1 at the stromal-immune interface. Candidate receptors for SVEP1 were identified by intersecting the interactome with the UniProt receptome (n = 1,656), resulting in 28 candidate receptor interactors (Fig. 3e, left), including integrins (ITGA9, ITGAM, ITGAX, ITGB5), adhesion receptors (ICAM1, CD93, COLEC12, ANTXR2, LRP1), and immune-modulatory receptors (CD55, ADGRG6, EGFR). Enrichment analysis (Fig. 3e, right; Fig. 3f) highlighted integrin signaling, ECM-cell interactions, complement cascades, leukocyte adhesion and migration, cytoskeletal remodeling, PI3K-Akt, and MAPK pathways. Analysis of receptor expression across human adipose cell populations (Fig. S5b–c) identified macrophage/monocyte, stromal/progenitor, and endothelial/stromal receptor modules. Macrophages exhibited a high proportion of SVEP1-associated receptors among immune cell subpopulations, with ITGA9 emerging as a particularly compelling candidate based on previous studies implicating SVEP1-ITGA9 interactions in macrophages. These findings support a model in which SVEP1 functions at the interface of ECM remodeling and myeloid cell communication.

**Figure 3.**
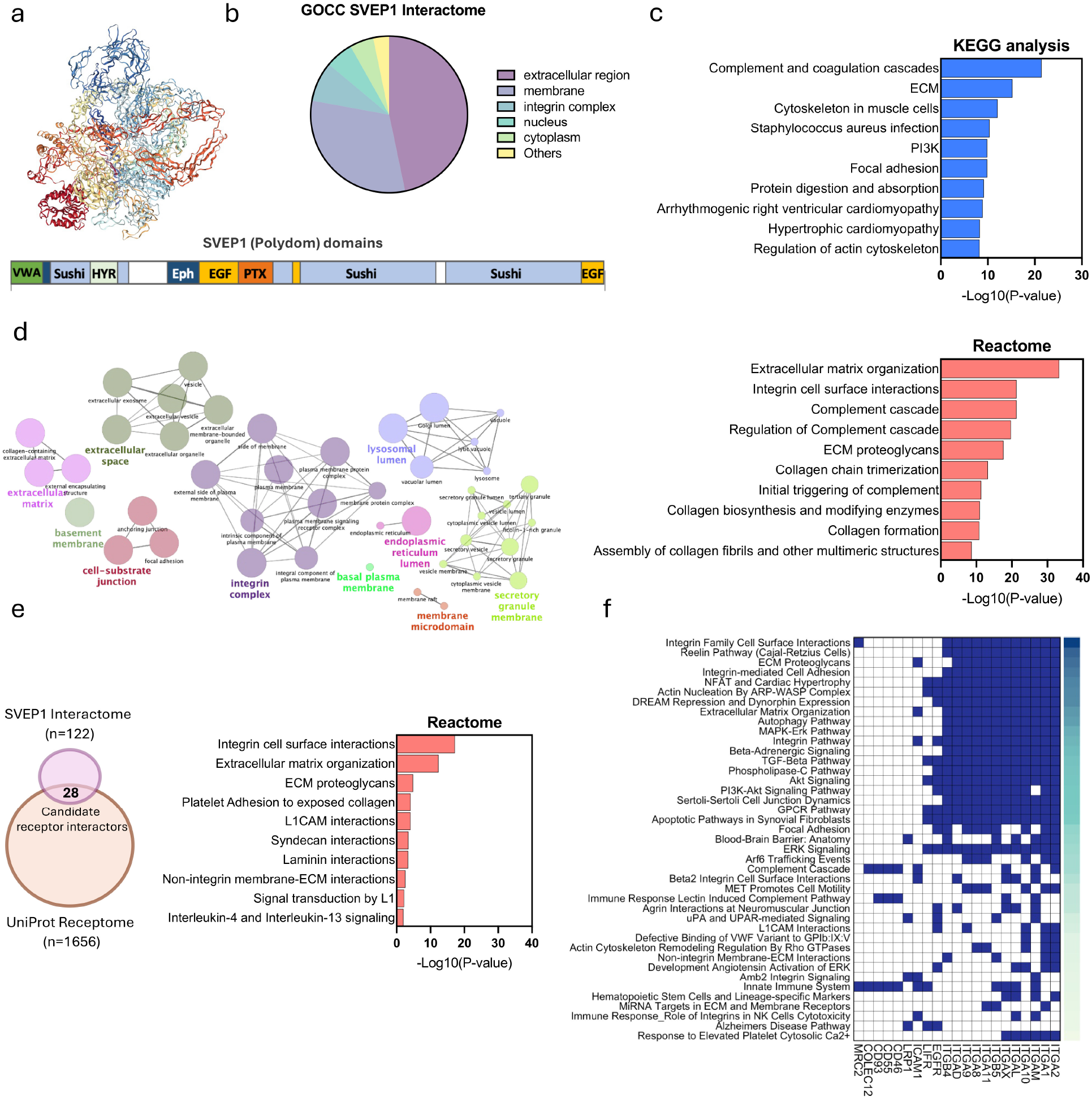
The predicted SVEP1 interactome links extracellular matrix organization with receptor-mediated immune signaling. (a) AlphaFold-predicted SVEP1 protein structure and its known domains. (b) GOCC classification of predicted SVEP1-interacting proteins (n = 122). (c) KEGG (top, blue) and Reactome (bottom, red) pathway enrichment analyses of the predicted SVEP1 interactome (FDR < 0.05). (d) Network visualization of enriched GOCC terms identified from the SVEP1 interactome using ClueGO. Functionally related terms are grouped based on shared protein membership; edges indicate term similarity derived from overlapping genes (right-sided hypergeometric test with Benjamini–Hochberg correction, FDR < 0.05). (e) Intersection of the SVEP1 interactome (n = 122) with the UniProt receptome (n = 1,656), identifying 28 candidate receptor interactors (left), followed by Reactome pathway enrichment analysis of the receptor-associated subset (right; FDR < 0.05). (f) Heatmap summarizing multi-source pathway associations of the SVEP1-associated receptors using GeneAnalytics, integrating pathway annotations from WikiPathways, GeneGo (MetaCore), Reactome, PubChem, QIAGEN, R&D Systems, and Cell Signaling Technology. Each tile denotes receptor participation in a given pathway

### *Svep1* deficiency alters glucose handling under chow diet conditions

To explore the metabolic role of SVEP1, we used heterozygous *Svep1*+/– mice, as homozygous deletion of *Svep1* is perinatally lethal. Six-week-old male *Svep1*+/+ and *Svep1*+/– littermates were maintained on a standard CHD for 12 weeks and subsequently subjected to metabolic and tissue analyses (Fig. 4a). qPCR analysis confirmed significantly reduced *Svep1* mRNA levels in inguinal white adipose tissue (iWAT) and epididymal white adipose tissue (eWAT), with a non-significant reduction in liver (Fig. 4b). Although organ-to-body weight ratios for iWAT, eWAT, BAT, and liver were comparable between the groups (Fig. 4c), *Svep1*+/– mice showed a trend toward lower body weight and a significant reduction in weight gain over time (Fig. 4d). Metabolic phenotyping indicated that *Svep1* deficiency alters glucose handling under basal chow conditions. Fasting blood glucose was lower in *Svep1*+/– mice after an overnight fast (Fig. 4e), while intraperitoneal glucose tolerance testing showed a trend toward improved glucose clearance (Fig. 4f). Insulin tolerance testing showed a comparable overall glucose excursion between genotypes (Fig. 4g). At the molecular level, qPCR analysis of eWAT revealed significant upregulation of *Pparg* and *Ppargc1a* in *Svep1*+/– animals (Fig. 4h), pointing to a shift in adipocyte metabolism and differentiation programming under basal nutritional conditions. Together, these data show that reduced *Svep1* dosage is sufficient to shift systemic glucose handling and adipose gene expression under standard nutritional conditions. Because *Svep1*+/– mice carry a constitutive, whole-body deletion, we next asked whether these changes originate in the adipocyte itself and turned to isolated adipose progenitors.

**Figure 4.**
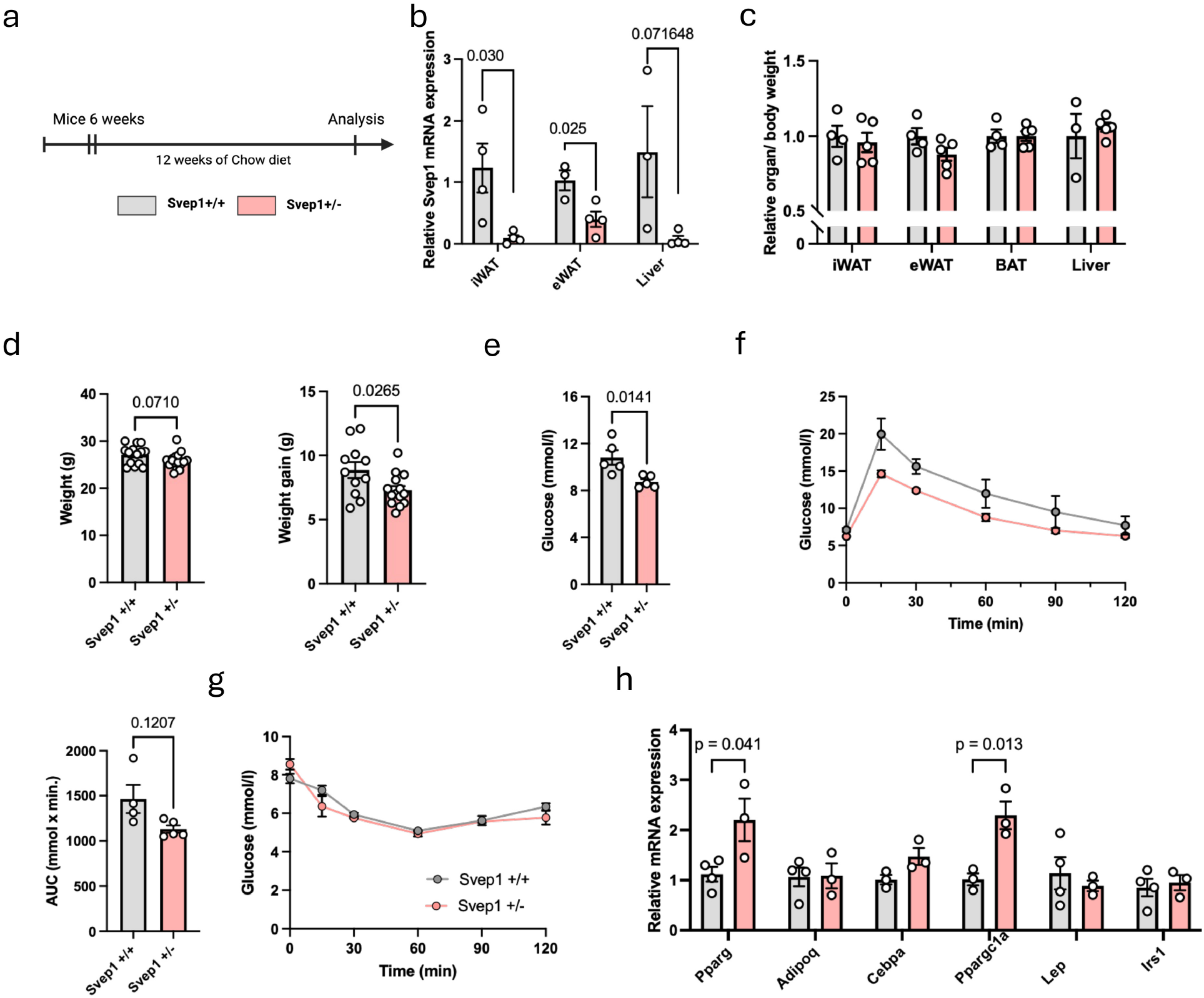
Metabolic consequences of *Svep1* deficiency (*Svep1*+/–) under chow diet conditions. (a) Experimental design. Six-week-old male *Svep1*+/+ and *Svep1*+/– mice were maintained on a standard chow diet for 12 weeks prior to metabolic and tissue analyses. (b) qPCR analysis of *Svep1* mRNA expression across metabolic tissues (iWAT, eWAT, and liver) in *Svep1*+/+ versus *Svep1*+/– mice (n = 4 per group). (c) Relative organ-to-body weight ratios for iWAT, eWAT, brown adipose tissue (BAT), and liver (n = 4–5 per group). (d) Total body weight and weight gain in *Svep1*+/+ and *Svep1*+/– mice (n = 11–13 per group). (e) Fasting blood glucose levels following an overnight fast (n = 5 per group). (f) Intraperitoneal GTT and corresponding AUC analysis (n = 5 per group); a non-significant trend toward improved glucose clearance was observed in *Svep1*+/– mice. (g) Intraperitoneal ITT analysis (n = 3 per group). (h) qPCR analysis of adipogenic and thermogenic gene expression in epididymal adipose tissue (n = 3–4 per group). Data are presented as mean ± SEM. Statistical significance for comparisons between two groups was assessed using two-tailed unpaired Student’s t-test or Mann–Whitney test, as appropriate. GTT and ITT curves were analyzed using two-way repeated-measures ANOVA.

### *Svep1* regulates adipocyte differentiation and metabolic function in vitro

To investigate the role of SVEP1 in adipocyte metabolism and adipogenesis, adipose-derived stromal/progenitor cells were isolated from iWAT of *Svep1*+/+ and *Svep1*+/– mice and differentiated into adipocytes in vitro (Fig. 5a). In wild-type cells, *Svep1* expression progressively increased during adipogenic differentiation (Fig. 5b), whereas differentiated *Svep1*+/– adipocytes exhibited significantly reduced *Svep1* mRNA compared with *Svep1*+/+ cells (Fig. 5c). Representative images and quantification after 14 days of differentiation (Fig. 5d) demonstrated that *Svep1*+/– cultures maintained adipogenic capacity, with a comparable LOA and similar bulk lipid accumulation between genotypes. Single-cell morphometric analysis demonstrated a significant reduction in lipid droplet (LD) area per cell in *Svep1*+/– adipocytes, while total LD number per cell and cell area remained unchanged. Overall lipid accumulation in these cultures was comparable despite the smaller droplet size, consistent with a higher number of adipocytes per field (Fig. 5e; Fig. S6a), indicating that *Svep1*+/– cultures contain more adipocytes with smaller lipid droplets while preserving lipid storage. At the molecular level, qPCR profiling (Fig. 5f) showed elevated transcripts of both *Pparg* and *Cd36* in *Svep1*+/– adipocytes, consistent with the eWAT findings observed in vivo. In the same cells, *Svep1*+/– adipocytes showed a significant increase in pAKT/tAKT upon insulin stimulation (Fig. 5g).

**Figure 5.**
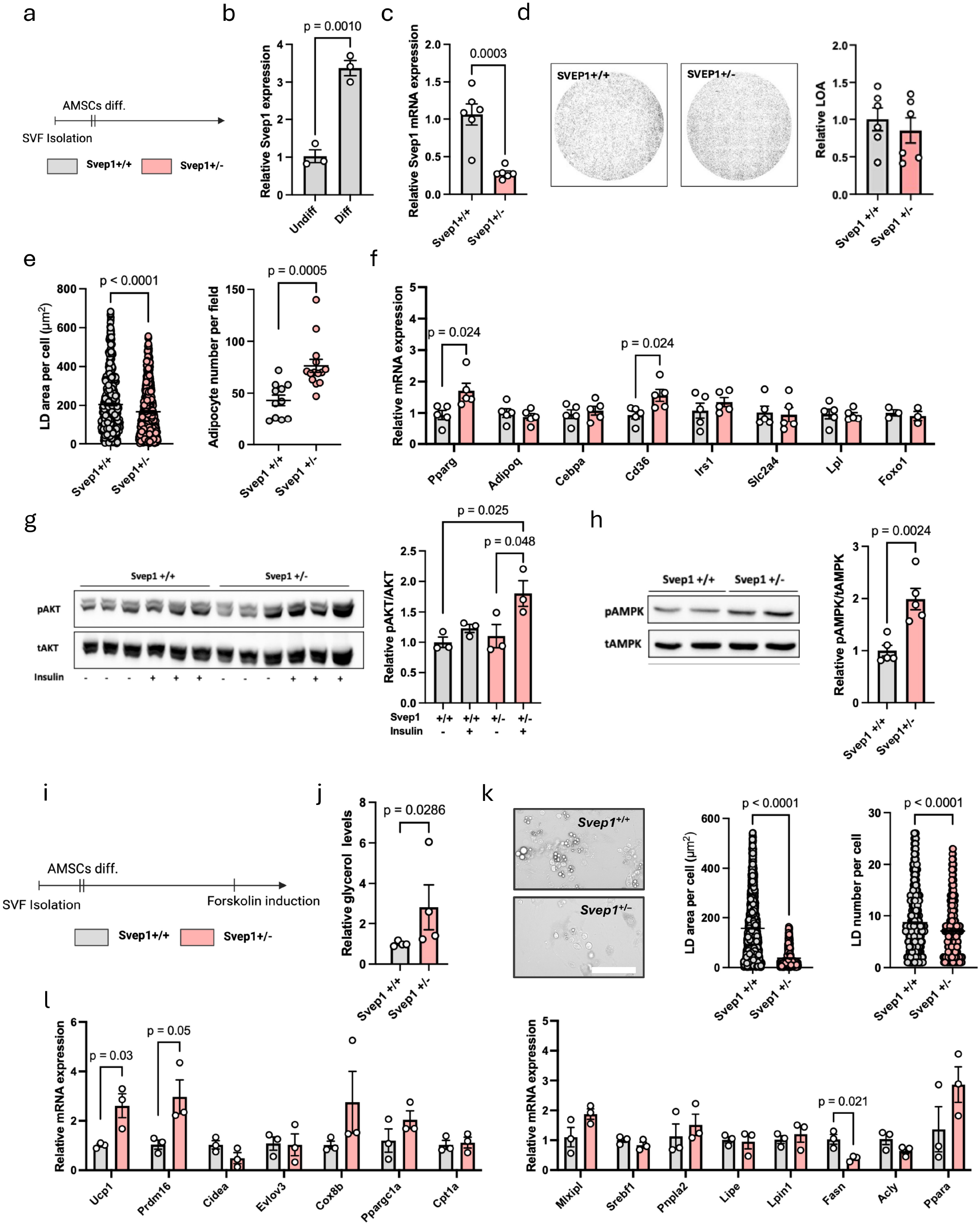
*Svep1* regulates adipocyte differentiation and metabolic function in vitro. (a) Experimental scheme. SVF cells were isolated from iWAT of *Svep1*+/+ and *Svep1*+/– mice and subjected to adipogenic differentiation in vitro. (b) qPCR analysis of *Svep1* expression in undifferentiated and differentiated adipose-derived mesenchymal stromal cells (n = 3). (c) Relative *Svep1* mRNA expression in differentiated adipocytes from *Svep1*+/+ and *Svep1*+/– cells (n = 6). (d) Representative images and quantification of adipogenic differentiation following 14 days of induction (n = 6). Scale bar = 100 µm. (e) Quantitative analysis of LD area per cell in *Svep1*+/+ (n = 543 cells from 11 cultures) and *Svep1*+/– (n = 540 cells from 14 cultures) adipocytes, and adipocyte number per field in *Svep1*+/+ (n = 11) and *Svep1*+/– (n = 14) cultures. (f) qPCR analysis of adipogenic and metabolic gene expression in differentiated adipocytes (n = 5–6). (g) Western blot analysis of pAKT and total AKT in differentiated *Svep1*+/+ and *Svep1*+/– adipose-derived mesenchymal stromal cells under basal (-) and insulin-stimulated (+) conditions, with quantification of the pAKT/tAKT ratio (n = 3). Significance was calculated using two-way ANOVA. (h) Western blot analysis of pAMPK and tAMPK, with quantification of the pAMPK/tAMPK ratio (n = 5). (i) Experimental scheme. SVF cells isolated from iWAT of *Svep1*+/+ and *Svep1*+/– mice were differentiated into adipocytes in vitro and subsequently treated with forskolin for 2 h. (j) Glycerol release into the culture medium after 2 h of forskolin stimulation (n = 4). (k) Representative images and quantification of LD area per cell and LD number per cell after 2 h of forskolin stimulation. Scale bar = 100 µm (n > 1,000 lipid droplets from 4 independent replicates). (l) qPCR analysis of metabolic and thermogenic gene expression in differentiated *Svep1*+/+ and *Svep1*+/– adipocytes following forskolin stimulation (n = 3). Data are presented as mean ± SEM. Statistical significance for comparisons between two groups was assessed using two-tailed unpaired Student’s t-test or Mann–Whitney test, as appropriate.

SVEP1 has been associated with adipocyte energy regulation, particularly lipid mobilization, and *Svep1* knockdown has been reported to enhance stimulated lipolysis and thermogenesis (*74*). Accordingly, AMPK activation was examined, revealing an elevated phosphorylated-to-total AMPK (pAMPK/tAMPK) ratio in *Svep1*+/– adipocytes (Fig. 5h). Given that AMPK activation promotes lipid mobilization, the lipolytic capacity of these cells was assessed directly. *Svep1*+/+ and *Svep1*+/– adipocytes were differentiated and stimulated with forskolin (Fig. 5i). Following forskolin stimulation, *Svep1*+/– adipocytes released more glycerol into the medium than *Svep1*+/+ controls (Fig. 5j). Morphometric analysis of the same cultures demonstrated a corresponding depletion of stored lipid; both LD area per cell and LD number per cell were substantially lower in *Svep1*+/– adipocytes (Fig. 5k). These findings indicate that *Svep1*-deficient adipocytes mobilize a greater proportion of their stored lipid upon forskolin stimulation. Post-stimulation qPCR profiling of metabolic and thermogenic transcripts (Fig. 5l) revealed increased *Ucp1* and *Prdm16* expression, along with reduced *Fasn* expression in *Svep1*+/– cells. Collectively, these in vitro data demonstrate that *Svep1* deficiency enhances the lipolytic capacity of adipocytes, accompanied by upregulation of thermogenic transcripts and downregulation of lipogenic genes.

### Prolonged fasting enhances AMPK activation and the catabolic response in *Svep1*-deficient mice

The pronounced lipolytic capacity of *Svep1*+/– adipocytes observed in vitro suggests that *Svep1* deficiency might also enhance fat mobilization in adipose tissue in vivo. To test whether this lipolytic phenotype is recapitulated at the tissue level and in whole-animal physiology, chow-fed *Svep1*+/+ and *Svep1*+/– mice were subjected to a prolonged fast (24–48 h; Fig. 6a). Following fasting, body weight was lower in *Svep1*+/– mice (Fig. 6b). Tissue weights of eWAT, iWAT, and liver across fed and fasted states (Fig. 6c) revealed genotype-specific differences, indicating an amplified response in *Svep1*+/– animals. This was accompanied by significantly lower fasting blood glucose levels in *Svep1*+/– mice (Fig. 6d) and improved glucose clearance, as indicated by a reduced area under the curve in the glucose tolerance test (Fig. 6e), in contrast to the modest trend observed under basal chow conditions (Fig. 4f). Western blot analysis of fasted epididymal adipose tissue (Fig. 6f) revealed a significantly higher pAMPK/tAMPK ratio in *Svep1*+/– mice, identifying AMPK as a candidate energy-sensing pathway through which *Svep1* deficiency amplifies the response to metabolic stress. Consistent with this signaling shift, qPCR analysis of fasted eWAT (Fig. 6g) revealed a coordinated shift away from lipogenesis and toward fatty-acid oxidation and lipid mobilization, with upregulation of *Ppara* and *Abhd5* alongside downregulation of the lipogenic regulators *Mlxipl* and *Srebf1* and a non-significant reduction in *Fasn*. Collectively, these data indicate that *Svep1* deficiency affects the adipose response to fasting, with activation of AMPK leading to improved glucose handling and a coordinated transcriptional response to nutrient demand.

**Figure 6.**
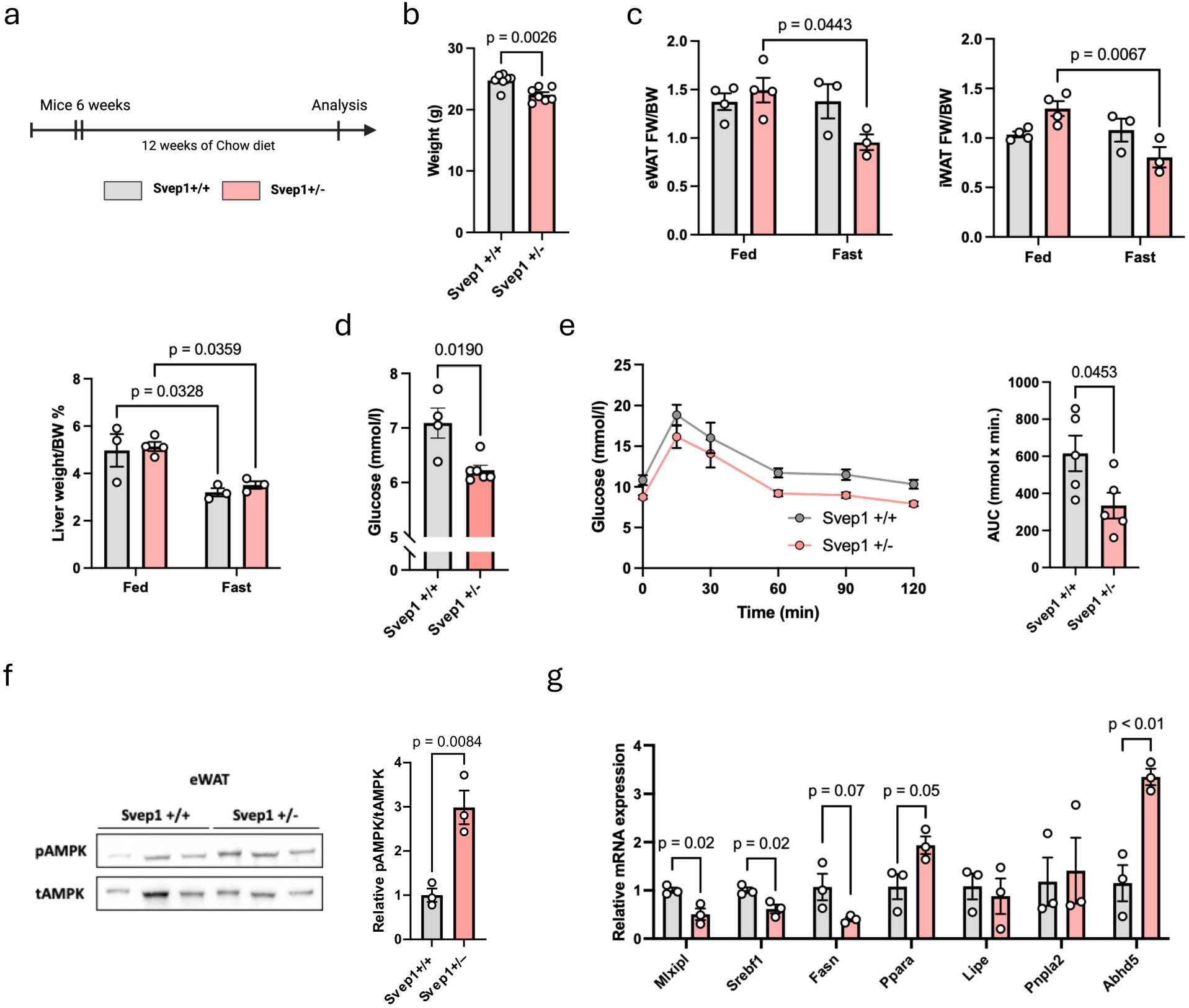
*Svep1* deficiency rewires the fasting response and amplifies AMPK signaling in adipose tissue. (a) Experimental design. Six-week-old male *Svep1*+/+ and *Svep1*+/– mice maintained on standard chow were subjected to a prolonged fast (24–48 h), with metabolic and tissue analyses performed at the end of the fasting period. (b) Body weight after the fast in fed versus fasted *Svep1*+/+ and *Svep1*+/– mice (n = 7 per group). (c) Tissue weights of eWAT, iWAT, and liver in fed and fasted *Svep1*+/+ and *Svep1*+/– mice, analyzed by two-way ANOVA (genotype × nutritional state) (n = 3–4 per group). (d) Fasting blood glucose levels measured at the end of the fast (n = 4–6). (e) Intraperitoneal GTT performed after the fast, with corresponding AUC analysis (n = 4–6 per group). (f) Western blot analysis of pAMPK and tAMPK in epididymal adipose tissue from fasted *Svep1*+/+ and *Svep1*+/– mice, with quantification of the pAMPK/tAMPK ratio (n = 3 per group). (g) qPCR analysis of metabolic gene expression in epididymal adipose tissue from fasted *Svep1*+/+ and *Svep1*+/– mice (n = 3 per group). Data are presented as mean ± SEM. Statistical significance for two-group comparisons was assessed using two-tailed unpaired Student’s t-test or Mann–Whitney test, as appropriate. Tissue weights in fed versus fasted conditions were analyzed by two-way ANOVA, and GTT curves by two-way repeated-measures ANOVA.

### *Svep1* deficiency alters systemic metabolism and adipose tissue remodeling under high-fat diet conditions

To determine whether the metabolic phenotype observed in *Svep1*+/– mice persists under chronic nutritional overload, *Svep1*+/+ and *Svep1*+/– littermates were subjected to a 60% HFD for 12 weeks, followed by metabolic and tissue analyses (Fig. 7a). Throughout the HFD period, *Svep1*+/– mice exhibited less weight gain compared with controls, resulting in lower final body weight and reduced longitudinal weight gain (Fig. 7b). After a 24 h fast, *Svep1*+/– mice lost a greater proportion of their initial body weight than *Svep1*+/+ controls (Fig. 7c), consistent with the enhanced fasting-induced lipid mobilization observed under chow conditions (Fig. 6). Organ-to-body weight ratio analysis indicated reduced iWAT and eWAT mass in HFD-fed *Svep1*+/– mice (Fig. 7d), suggesting attenuated adipose expansion. Following an overnight fast, *Svep1*+/– mice had significantly lower fasting blood glucose levels than *Svep1*+/+ controls (Fig. 7e). Intraperitoneal glucose tolerance testing showed improved glucose clearance in *Svep1*+/– mice (Fig. 7f), and insulin tolerance testing demonstrated enhanced insulin responsiveness, indicating improved systemic insulin sensitivity (Fig. 7g). Whole-mount imaging of visceral adipose tissue revealed markedly smaller adipocytes in HFD-fed *Svep1*+/– mice, with significantly reduced mean lipid droplet size in epididymal adipose tissue (Fig. 7h). qPCR analysis of adipogenic and metabolic gene expression in eWAT and iWAT (Fig. 7i) showed elevated *Pparg* and *Cebpa* transcripts in eWAT and increased *Pparg* in iWAT of HFD-fed *Svep1*+/– mice, indicating preserved or enhanced adipogenic programming despite reduced adipocyte size. Hepatic qPCR analysis (Fig. S7a) further demonstrated marked upregulation of *Fgf21* in *Svep1*+/– mice. Collectively, these findings indicate that *Svep1* deficiency confers a metabolically favorable phenotype under HFD-induced obesity, characterized by attenuated adipose tissue expansion, reduced adipocyte hypertrophy, and improved systemic glucose handling and insulin sensitivity.

**Figure 7.**
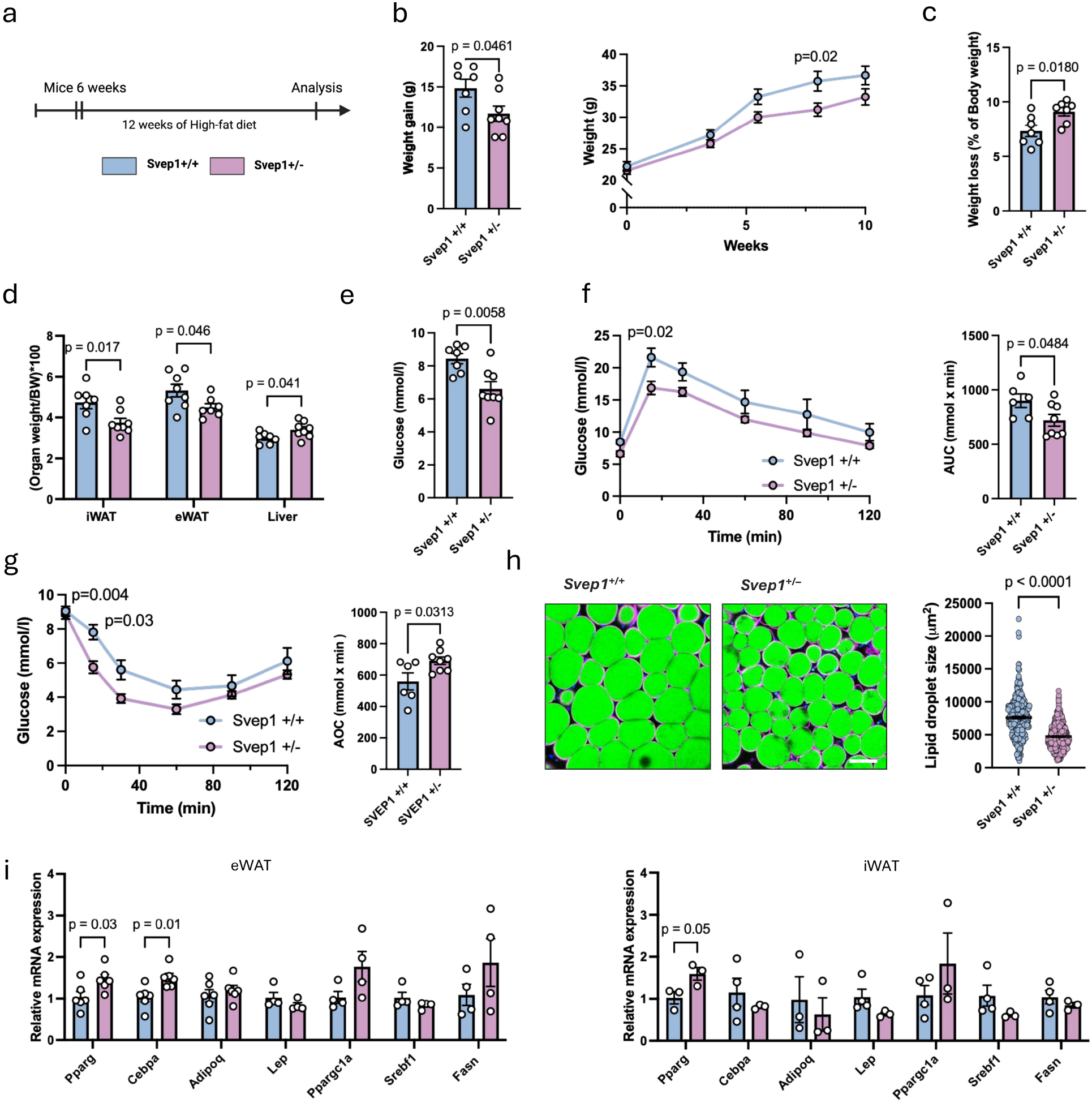
*Svep1* deficiency alters systemic metabolism and adipose tissue remodeling under high-fat diet conditions. (a) Experimental design. Six-week-old male *Svep1*+/+ and *Svep1*+/– mice were fed a HFD (60% kcal from fat) for 12 weeks, followed by metabolic and tissue analyses. (b) Body weight gain at endpoint and longitudinal body weight during HFD feeding (n = 7–8 per group). (c) Body weight loss following a 24 h fast, expressed as percentage of initial body weight (n = 7–8 per group). (d) Relative organ-to-body weight ratios for iWAT, eWAT, and liver (n = 7–8 per group). (e) Fasting blood glucose levels following a 12 h fast (n = 7–8 per group). (f) Intraperitoneal GTT with corresponding AUC analysis (n = 6–8 per group). (g) Intraperitoneal ITT with corresponding area over the curve (AOC) analysis (n = 6–8 per group). (h) Representative whole-mount images of visceral adipose tissue with quantification of lipid droplet size (representative measurements pooled from 3 mice per group, >200 droplets per mouse). Scale bar = 100 µm. (i) qPCR analysis of adipogenic and metabolic gene expression in epididymal and subcutaneous adipose tissue (n = 3–6). Data are presented as mean ± SEM. Statistical significance for comparisons between two groups was assessed using two-tailed unpaired Student’s t-test or Mann–Whitney test, as appropriate. GTT and ITT curves were analyzed using two-way repeated-measures ANOVA.

To investigate how reduced *Svep1* expression alters the transcriptional program of adipose tissue during HFD-induced obesity, bulk RNA-seq was performed on eWAT from HFD-fed *Svep1*+/+ and *Svep1*+/– mice. Differential expression analysis identified 159 upregulated and 247 downregulated genes in *Svep1*+/– adipose tissue (adjusted p < 0.05; |log₂FC| ≥ 0.5; Fig. 8a). GOBP analysis (Fig. 8b) indicated that upregulated pathways were associated with the tricarboxylic acid cycle and central carbon metabolism, fatty acid and lipid synthesis, carbohydrate and glucose homeostasis, and thermogenesis, including increased expression of lipogenic and metabolic genes (*Acaca*, *Acly*, *Srebf1*). The adipogenic regulator *Pparg* was also elevated, consistent with enhanced adipocyte differentiation, insulin responsiveness, and metabolic activity. In contrast, downregulated pathways were predominantly linked to myeloid/leukocyte activation and trafficking, phagocytosis, and ECM/collagen remodeling, with significant decreases in myeloid activation markers (*Itgax*, *Adgre1*, *Trem2*, *Ptprc*), macrophage surface receptors (*Cd68*, *Cd84*), and ECM transcripts such as collagen genes (*Col12a1*, *Col16a1*), matrix metalloproteinases, and cathepsins. Multi-source GeneAnalytics pathway integration (Fig. 8c) confirmed that downregulated programs were centered on immune activation, antigen presentation, cytokine signaling, and ECM-cell interaction modules. In contrast, upregulated programs were enriched in general metabolism, glycolysis, adipogenesis, glucose and energy metabolism, and cholesterol/lipid homeostasis (SREBF/MiR33 pathway). PROGENy pathway modeling (Fig. 8d; Fig. S8a) demonstrated decreased activity of NF-κB, TNFα, and TGF-β signaling pathways in *Svep1*+/– adipose tissue, with PI3K and TGF-β pathways identified as the most strongly altered. Transcription factor activity analysis using DoRothEA (Fig. 8e; Fig. S8b) further revealed decreased activity of transcription factors associated with myeloid activation and ECM remodeling, including SPI1, RELA, and NFKB1, alongside increased activity of transcription factors linked to lipid handling and metabolic regulation, such as THRB, SREBF1, and DBP.

**Figure 8.**
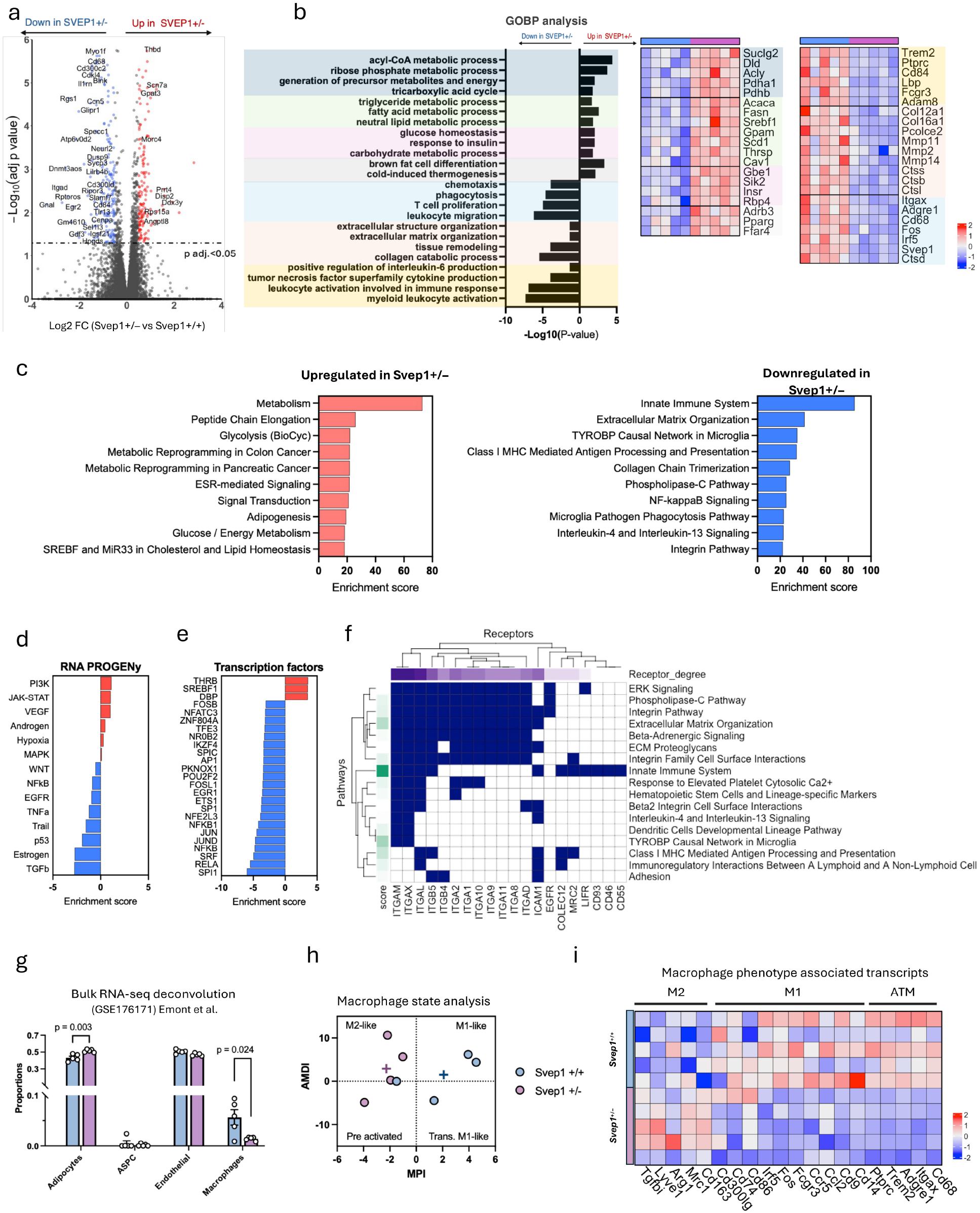
*Svep1* deficiency drives immunometabolic transcriptional reprogramming in adipose tissue under high-fat diet. (a) Volcano plot of DEGs in epididymal adipose tissue from HFD-fed *Svep1*+/+ and *Svep1*+/– mice (n = 5 per group; adjusted p < 0.05; |log₂FC| ≥ 0.5), highlighting significantly upregulated (red) and downregulated (blue) genes. (b) GOBP enrichment analysis of DEGs (FDR < 0.05; |log₂FC| ≥ 0.5), with representative heatmaps of genes contributing to selected upregulated and downregulated pathways (n = 5 per group). (c) Pathway enrichment analysis of DEGs using GeneAnalytics, showing top upregulated (left) and downregulated (right) pathways. (d) PROGENy pathway activity analysis estimating differential signaling pathway activity between genotypes. (e) Transcription factor activity inference using decoupleR/DoRothEA, highlighting the top differentially active transcription factors between *Svep1*+/+ and *Svep1*+/– mice. (f) Heatmap integrating downregulated DEG-derived pathways with SVEP1-associated receptor-linked pathways, highlighting shared signaling modules between transcriptional changes and receptor-level interactions. (g) Bulk RNA-seq cell-type deconvolution using the Emont et al. (GSE176171) adipose single-cell dataset, showing estimated proportions of adipocytes, ASPCs, endothelial cells, and macrophages (n = 5 per group). (h) MPI and AMDI analysis using MacSpectrum, indicating shifts in macrophage activation states (n = 4 per group). (i) Heatmap of macrophage phenotype-associated transcripts, including M1-like, M2-like, and ATM markers (n = 5 per group). Data are presented as mean ± SEM. Statistical significance for comparisons between two groups was assessed using two-tailed unpaired Student’s t-test or Mann–Whitney test, as appropriate.

To determine whether the altered transcriptional landscape of *Svep1*+/– adipose tissue corresponds to receptor-linked signaling networks, the previously identified SVEP1-associated receptor set (Fig. 3) was examined. A heatmap integrating downregulated DEG-derived pathways with SVEP1-associated receptor-linked pathways (Fig. 8f) demonstrated that a substantial proportion of suppressed signaling modules corresponded to pathways associated with SVEP1 receptors, including integrin-mediated adhesion, complement cascades, and inflammatory signaling involving MAPK families. BulkSignalR ligand-receptor modeling (Fig. S8c) further indicated a broad reduction in ligand-receptor interactions related to integrin signaling, chemokine-mediated communication, and ECM-immune crosstalk in *Svep1*+/– adipose tissue. Weighted receptor family and receptor-level involvement analysis (Fig. S8d–e) confirmed that integrin receptors represented the dominant share of receptor-pathway intersections, with network representations illustrating the functional connections between SVEP1-associated receptor nodes and suppressed downstream pathways. Deconvolution of bulk RNA-seq data using adipose tissue snRNA-seq reference profiles (Fig. 8g) revealed a lower estimated proportion of macrophages and a relative increase in adipocyte representation in *Svep1*+/– adipose tissue, providing transcriptome-level evidence of reduced immune infiltration. MacSpectrum analysis of the MPI and AMDI (Fig. 8h) indicated that *Svep1*+/– mice exhibited a shift from a pro-inflammatory M1-like macrophage state toward an alternatively activated, M2-like profile. A heatmap of representative macrophage phenotype-associated transcripts (Fig. 8i) supported these findings, with higher M2-associated markers (*Mrc1*, *Arg1*) in *Svep1*+/– tissue and more prominent M1 markers (*Cd74*, *Cd300*, *Cd68*) in *Svep1*+/+ samples. Collectively, these results suggest that reduced *Svep1* expression shifts obese adipose tissue away from an ECM-and inflammatory-activated state toward a more metabolically favorable state, implicating integrin-centered receptor signaling as a key interface through which SVEP1 may regulate macrophage abundance and phenotype.

### SVEP1 modulates adipocyte-macrophage interactions and integrin-dependent macrophage functional responses

The convergence of transcriptomic, pathway, and receptor analyses in macrophages prompted direct characterization of this compartment in HFD-fed *Svep1*+/+ and *Svep1*+/– adipose tissue. Whole-mount immunofluorescence staining of CD11b in visceral adipose tissue (Fig. 9a) revealed significantly fewer CD11b+ cells per field in *Svep1*+/– animals, a finding confirmed by flow cytometry of the stromal vascular fraction (Fig. 9b). The reduction in macrophage abundance was attributable to a specific decrease in the CD11c+ pro-inflammatory ATM subset, consistent with a shift away from classically activated, obesity-associated macrophages. Because ITGA9 emerged as the leading candidate SVEP1 receptor on adipose macrophages in the interactome analysis (Fig. 3e; Fig. S5b–c), we also tested whether the predicted SVEP1-ITGA9 axis is reflected in vivo. ITGA9 mean fluorescence intensity was quantified on CD45+CD11b+F4/80+ macrophages from the same animals (Fig. 9c). ITGA9 levels were reduced in *Svep1*+/– macrophages, suggesting an ITGA9-mediated macrophage-ECM interaction related to *Svep1*. qPCR profiling of macrophage-associated transcripts (Fig. 9d) demonstrated reduced *Ccl2* and *Tnfa* expression in *Svep1*+/– adipose tissue, supporting a less pro-inflammatory state of the adipose macrophage population. Together with the transcriptomic and phenotypic shifts characterized in Fig. 8, these data indicate that *Svep1* deficiency modulates the adipose macrophage compartment by reducing both macrophage abundance and inflammatory tone. To further dissect adipocyte-macrophage crosstalk under inflammatory conditions, an in vitro co-culture of *Svep1*+/– adipocytes with wild-type BMDMs was challenged with lipopolysaccharide (LPS) for 8 h (Fig. 9e). These co-cultures (Fig. 9f) exhibited both metabolic reprogramming and reduced inflammation. In contrast, LPS stimulation of *Svep1*+/+ and *Svep1*+/– adipocytes alone produced no inflammatory differences between genotypes (Fig. S9a). Collectively, these results indicate that the observed phenotype requires the adipocyte-macrophage interaction.

**Figure 9.**
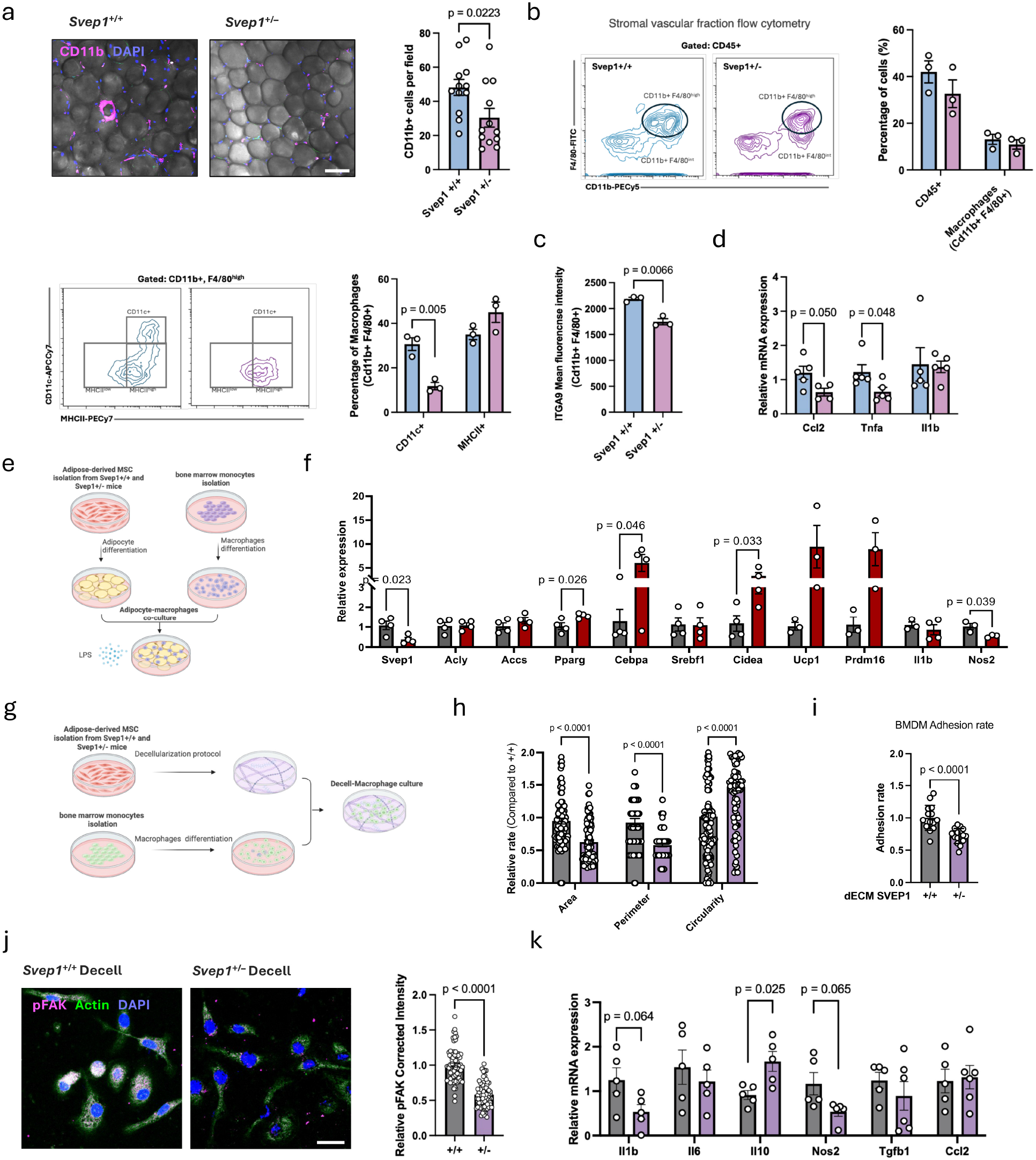
SVEP1 regulates adipocyte-macrophage crosstalk and macrophage functional responses. (a) Representative immunofluorescence images of visceral adipose tissue stained for CD11b (magenta) and DAPI (blue) from *Svep1*+/+ and *Svep1*+/– mice, with quantification of CD11b+ cells per field. Each dot represents one quantified field (12 per group); quantification was performed across three biological replicates per group. Scale bar = 100 µm. (b) Flow cytometry analysis of the SVF gated on CD45+ immune cells. Representative dot plots show CD11b and F4/80 staining in *Svep1*+/+ and *Svep1*+/– mice (top left), with quantification of total CD45+ cells and macrophages (CD11b+F4/80+) (top right), and analysis of the F4/80^high^ ATM population. Representative contour plots of MHCII versus CD11c staining (bottom left), with quantification of CD11c+ and MHCII+ macrophage subsets (bottom right) (n = 3). (c) Mean fluorescence intensity of ITGA9 in macrophages (CD45+CD11b+F4/80+) isolated from HFD-fed *Svep1*+/+ and *Svep1*+/– mice (n = 3). (d) qPCR analysis of macrophage-associated gene expression in HFD-fed *Svep1*+/+ and *Svep1*+/– mice (n = 4–5). (e) Experimental scheme for the adipocyte-macrophage co-culture system. AMSCs from *Svep1*+/+ and *Svep1*+/– mice were differentiated into adipocytes, co-cultured with bone marrow-derived macrophages, and stimulated with LPS for 8 h. (f) qPCR analysis of adipogenic, metabolic, thermogenic, and inflammatory gene expression in adipocyte-macrophage co-cultures (n = 4). (g) Experimental scheme for the dECM-macrophage culture system. AMSCs from *Svep1*+/+ and *Svep1*+/– mice were used to generate decellularized matrices, followed by culture of bone marrow-derived macrophages on the resulting ECM. (h) Quantification of macrophage morphological parameters, including cell area, perimeter, and circularity, in bone marrow-derived macrophages cultured on *Svep1*+/+-or *Svep1*+/–-derived decellularized matrices (n = 108 and 94 cells, respectively, from 4–5 independent replicates). (i) Adhesion of wild-type bone marrow-derived macrophages seeded onto decellularized extracellular matrices generated from *Svep1*+/+ or *Svep1*+/– AMSCs (n = 15 fields, from 3 independent replicates). (j) Immunofluorescence analysis and quantification of relative pFAK corrected total cell fluorescence (CTCF) in bone marrow-derived macrophages cultured on *Svep1*+/+ (n = 83 cells) or *Svep1*+/– (n = 73 cells)-derived decellularized matrices, from 4 independent replicates. Scale bar = 100 µm. (k) qPCR analysis of inflammatory gene expression in bone marrow-derived macrophages cultured on decellularized matrices (n = 5). Data are presented as mean ± SEM. Statistical significance for comparisons between two groups was assessed using two-tailed unpaired Student’s t-test or Mann–Whitney test, as appropriate.

To isolate the contribution of SVEP1-dependent matrix composition from cellular signals, dECM was generated from *Svep1*+/+ and *Svep1*+/– AMSCs, and BMDMs were seeded onto these cell-free matrices (Fig. 9g). Macrophages cultured on *Svep1*+/– matrices exhibited altered cell area, perimeter, and circularity (Fig. 9h), as well as altered adhesion compared to those on *Svep1*+/+ matrices (Fig. 9i). These findings demonstrate that matrix-intrinsic properties are sufficient to reshape macrophage morphology and attachment. Immunofluorescence quantification of phosphorylated focal adhesion kinase (pFAK) corrected total cell fluorescence (CTCF; Fig. 9j) showed significantly reduced FAK phosphorylation in macrophages on *Svep1*+/– matrices, indicating that SVEP1 engages integrin-FAK signaling in these cells. Consistent with this signaling shift, qPCR profiling of BMDMs cultured on these matrices (Fig. 9k) revealed trends toward reduced *Il1b* and *Nos2* expression, alongside a significant increase in *Il10*.

In summary, these data identify SVEP1 as an extracellular matrix-anchored regulator of adipose tissue macrophages. *Svep1* deficiency reduces macrophage accumulation in vivo, shifts ATM phenotypes away from pro-inflammatory states, and attenuates inflammatory cytokine and chemokine expression. Collectively, these findings position SVEP1 as a contributor to the maintenance of the inflammatory adipose niche during diet-induced obesity.

## Discussion

In this study, we identify SVEP1 as a component of the adipose niche, whose expression is upregulated in visceral fat under an HFD. By integrating bulk RNA-seq and LC-MS/MS proteomics with pathway inference, transcription factor activity profiling, and ligand-receptor analyses, we delineate an ECM-centered signaling axis in which SVEP1 sustains an adhesion-and inflammation-permissive tissue niche. Characterization across human datasets, including single-cell transcriptomics and publicly available clinical cohort data, confirmed that *SVEP1* is preferentially expressed in adipocytes and positively associated with markers of adiposity, insulin resistance, and systemic inflammation. Construction of the SVEP1 interactome and receptome revealed that macrophages harbor the highest density of SVEP1-associated receptors, with integrins as the dominant receptor family. Consistent with this model, *Svep1* deficiency conferred protection during diet-induced obesity: adipose tissue exhibited reduced adipocyte hypertrophy, attenuated pro-inflammatory macrophage accumulation, and a shift toward oxidative and thermogenic metabolic programs, accompanied by improved systemic glucose levels and clearance. Functional assays using adipocyte-macrophage co-cultures and decellularized matrices further demonstrated that SVEP1-dependent ECM directly regulates macrophage adhesion, morphology, polarization, and integrin signaling.

Our integrative multi-omics analyses showed that ECM expansion and adhesion-related signaling constitute major biological programs induced by HFD in visceral adipose tissue (*5*, *8*). Transcriptomic profiling showed coordinated upregulation of ECM organization, collagen biosynthesis, and cell-matrix adhesion pathways, accompanied by suppression of oxidative phosphorylation and mitochondrial functions. Proteomic data from the same depots recapitulated these results, with increased abundance of extracellular and basement-membrane proteins and concurrent reductions in mitochondrial and metabolic proteins. Several ECM components with established roles in adipose remodeling were among the most strongly induced. This included collagen VI, whose depletion reduces adipose fibrosis and improves insulin sensitivity (*75*), and fibrillin-1, an elastic-fiber constituent linked to adipocyte size and TGF-β-dependent remodeling (*76*). Focusing the analysis specifically on the ECM compartment, SVEP1 consistently ranked among the most prominently and concordantly induced matrix components across both RNA and protein modalities, nominating it as a strong candidate mediator of obesity-driven adipose remodeling.

SVEP1 is a multidomain protein containing VWA, CCP/Sushi, EGF, and pentraxin motifs that localizes to the adipocyte extracellular matrix and basement-membrane compartments. It functions as a ligand for integrin α9β1 (*16*), PEAR1 (*18*), and Tie1 (*19*), and activates downstream MAPK-ERK, AKT-mTOR, and NF-κB signaling in vascular and immune contexts (*20*, *25*). Most of what is known about SVEP1 function comes from the vasculature. Homozygous loss of *Svep1* in mice results in lymphatic vessel defects and perinatal lethality, indicating an evolutionarily conserved requirement in vascular integrity (*77*). Human genetic studies link *SVEP1* coding variants and elevated circulating levels to coronary artery disease, hypertension, and type 2 diabetes (*21*, *22*), positioning SVEP1 as an ECM-embedded signaling regulator with established functions in mesenchymal-vascular communication.

Within adipose tissue, SVEP1 has been detected in matrisome studies as part of the adipocyte ECM and secreted protein repertoire. In transcriptomic analyses, SVEP1 was reported to be enriched in subcutaneous and visceral fat depots of individuals with obesity (*23*, *24*). In our study, SVEP1 protein abundance was significantly higher in epididymal white adipose tissue of HFD-fed mice. Human meta-analysis data confirmed a positive correlation between *SVEP1* transcript levels and BMI (*51*). Furthermore, a study utilizing a 236-subject subcutaneous adipose tissue RNA-seq dataset found that *SVEP1* expression correlated positively with BMI, fat mass, fat percentage, and waist circumference, supporting the clinical relevance of SVEP1 as an obesity-associated ECM component (*74*). At single-cell resolution, *SVEP1* expression is primarily localized to the adipocyte lineage, particularly ASPCs and mature adipocytes (*43*). Collectively, these datasets establish that SVEP1 is produced within the adipocyte lineage and that its expression scales with adiposity.

Emerging data indicate that SVEP1 participates in immune regulation. *SVEP1* transcription is responsive to inflammatory stimuli including TNFα, and its promoter contains an NF-κB binding site (*29*, *78*). Several studies have identified SVEP1 as a sepsis-associated gene, with knockdown models showing altered pro-inflammatory cytokine expression (*79*, *80*), and in an atherosclerosis model, smooth muscle-specific *Svep1* deletion alters cytokine production and macrophage infiltration (*20*, *78*, *81*). Building on these observations, we show here, using a genetic *Svep1*+/– model, that this immune role also operates within the adipose niche. MacSpectrum analysis of both HFD-fed wild-type and *Svep1*+/– adipose tissue demonstrated that reduced SVEP1 levels shift macrophage polarization away from a pro-inflammatory M1-like state, with lower MPI values and increased expression of M2-associated transcripts such as *Mrc1* and *Arg1*. Flow cytometry confirmed a selective reduction in CD11c+ pro-inflammatory macrophages in *Svep1*+/– adipose tissue under HFD. These findings place SVEP1 within an ECM-linked innate immune signaling axis in which matrix composition directly influences macrophage polarization. This study also links the ECM to adipocyte energy metabolism: reducing SVEP1, a protein produced by the adipocyte lineage itself, enhances the metabolic and lipolytic capacity of these cells without impairing differentiation. *Svep1*+/– adipocytes display increased expression of adipogenic transcripts (*Pparg*), a significant insulin-stimulated increase in AKT phosphorylation, activated AMPK signaling, and elevated forskolin-stimulated lipolysis. Previous studies demonstrated that shRNA-mediated *Svep1* knockdown enhances stimulated lipolysis in white adipocytes without affecting adipogenic differentiation (*74*), and the *Svep1*+/– model recapitulates this lipolytic phenotype. Collectively, these results indicate an intrinsic role for SVEP1 in adipocytes and, together with the fasting and HFD phenotypes, imply that its loss may modestly promote adipocyte energy sensing, lipid mobilization, and insulin responsiveness. These findings are consistent with broader evidence that ECM glycoproteins can directly regulate adipocyte energy metabolism: LAMA4 silencing enhances AMPK-PGC1α signaling and metabolic gene expression, while MAGP1 deficiency impairs cold-induced browning (*82*, *83*). Mechanistically, the composition of the ECM determines its mechanical properties, which resident cells convert into metabolic output (*5*, *8*). In HFD-fed mice, visceral adipose ECM loses elastin and accumulates collagenous proteins, resulting in a stiffer niche. This stiffness is associated with increased inhibitory serine phosphorylation of IRS-1, reduced plasma-membrane GLUT4, and diminished insulin sensitivity (*84*). Experimentally increasing the stiffness of the adipocyte niche in culture reproduces this pattern, decreasing AKT phosphorylation and suppressing adipogenic gene expression (*85*). Conversely, defined biomechanical stimulation alone is sufficient to remodel the adipocyte proteome and increase mitochondrial activity and lipid-droplet content (*86*). The observation that *Svep1* deficiency de-represses adipocyte energy-handling programs suggests that it may function as an ECM-based regulator of adipose energy homeostasis.

Our data extend the current understanding of SVEP1 within adipose immune metabolism by establishing a multi-layered model that links ECM composition to both immune and metabolic outcomes. The construction of the SVEP1 interactome and receptome constitutes a novel contribution, revealing that SVEP1-associated receptors are enriched in integrin, complement, and adhesion families. Receptor-pathway intersection analysis demonstrated that transcriptional changes in *Svep1*+/– adipose tissue under HFD conditions map directly onto signaling modules connected to these receptors, with integrins accounting for the majority of receptor-pathway intersections. Functional validation using decellularized ECM and co-culture assays confirmed that SVEP1-dependent matrix cues are sufficient to alter macrophage adhesion, morphology, inflammatory gene expression, and integrin-FAK phosphorylation. Collectively, these data support a model in which SVEP1 plays a role in an ECM-integrin-macrophage signaling axis. Under HFD, increased SVEP1 deposition strengthens integrin-mediated adhesion, promotes pro-inflammatory macrophage retention and activation, and concurrently restrains adipocyte metabolic capacity.

In contrast, reduced SVEP1 alters this program, resulting in coordinated attenuation of immune activation and enhancement of thermogenic and lipolytic function. Although the receptome analysis identified integrin α9β1 as the most prominently macrophage-enriched integrin in the SVEP1 protein landscape, the functional assays in this study addressed the broader ECM-integrin axis.

In summary, our findings identify SVEP1 as an ECM-anchored regulator that couples structural remodeling to macrophage activation and metabolic restraint during nutritional overload, highlighting SVEP1 and its receptor network as potential targets for modulating adipose tissue immuno-metabolic function in obesity.

## Acknowledgments

This work was performed in partial fulfillment of the requirements for the PhD degree of Nadav Kislev, Faculty of Medicine. Nadav Kislev is an awardee of the Marian Gertner Institute for Medical Nanosystems, Tel Aviv University Center for Combating Pandemics (TCCP), and the Healthy Longevity Research Center, Tel Aviv University.

## Author contributions

N.Ki.: conceptualization, methodology, investigation, formal analysis, bioinformatics analysis, data curation, visualization, original draft writing. N.Ka. and R.I.: conceptualization support and experimental handling. D.B.: conceptualization, supervision, funding acquisition, review & editing. All authors critically reviewed the manuscript and approved the final version for publication.

## Data and materials availability

All data associated with this study are present in the paper or the Supplementary Materials, and any additional data will be provided by the corresponding author upon reasonable request. All analyses used the publicly available software packages listed in Methods.

## Supplementary Figures

**Figure S1.**
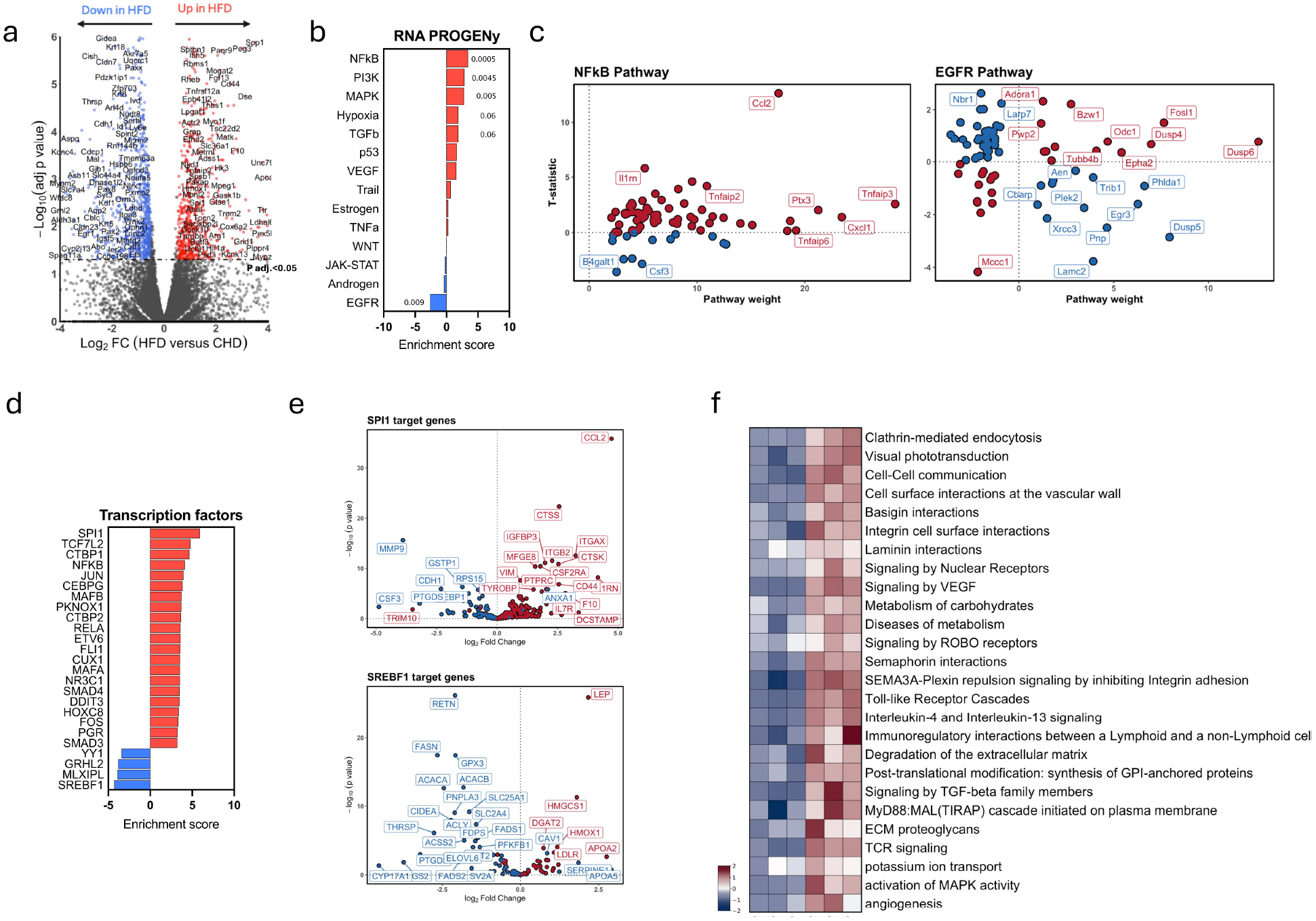
Pathway-, transcription factor-, ligand-receptor-, and ECM-focused transcriptomic analyses of visceral adipose tissue in HFD versus CHD. (a)Volcano plot of DEGs in epididymal adipose tissue from HFD-and CHD-fed mice, showing log₂ fold change versus adjusted p value and highlighting significantly upregulated (red) and downregulated (blue) genes. (b) PROGENy pathway activity analysis showing major signaling pathways altered in HFD versus CHD visceral adipose tissue, with p values indicated. (c) PROGENy contribution plots highlighting genes contributing most strongly to the top upregulated and top downregulated pathways (NF-κB pathway, top; EGFR pathway, bottom) in HFD versus CHD visceral adipose tissue. Red indicates genes contributing positively to pathway activation, and blue indicates genes contributing negatively. (d) Transcription factor activity analysis using decoupleR/DoRothEA showing the top differentially active transcription factors between HFD and CHD samples. (e) Volcano plots of transcription factor target genes corresponding to the most differentially active transcription factors identified by decoupleR/DoRothEA between HFD and CHD samples (SPI1 target genes, top; SREBF1 target genes, bottom). (f) BulkSignalR ligand-receptor pathway analysis comparing HFD and CHD samples, showing pathway-level regulation inferred from ligand-receptor interactions and highlighting signaling programs enriched or suppressed under HFD conditions

**Figure S2.**
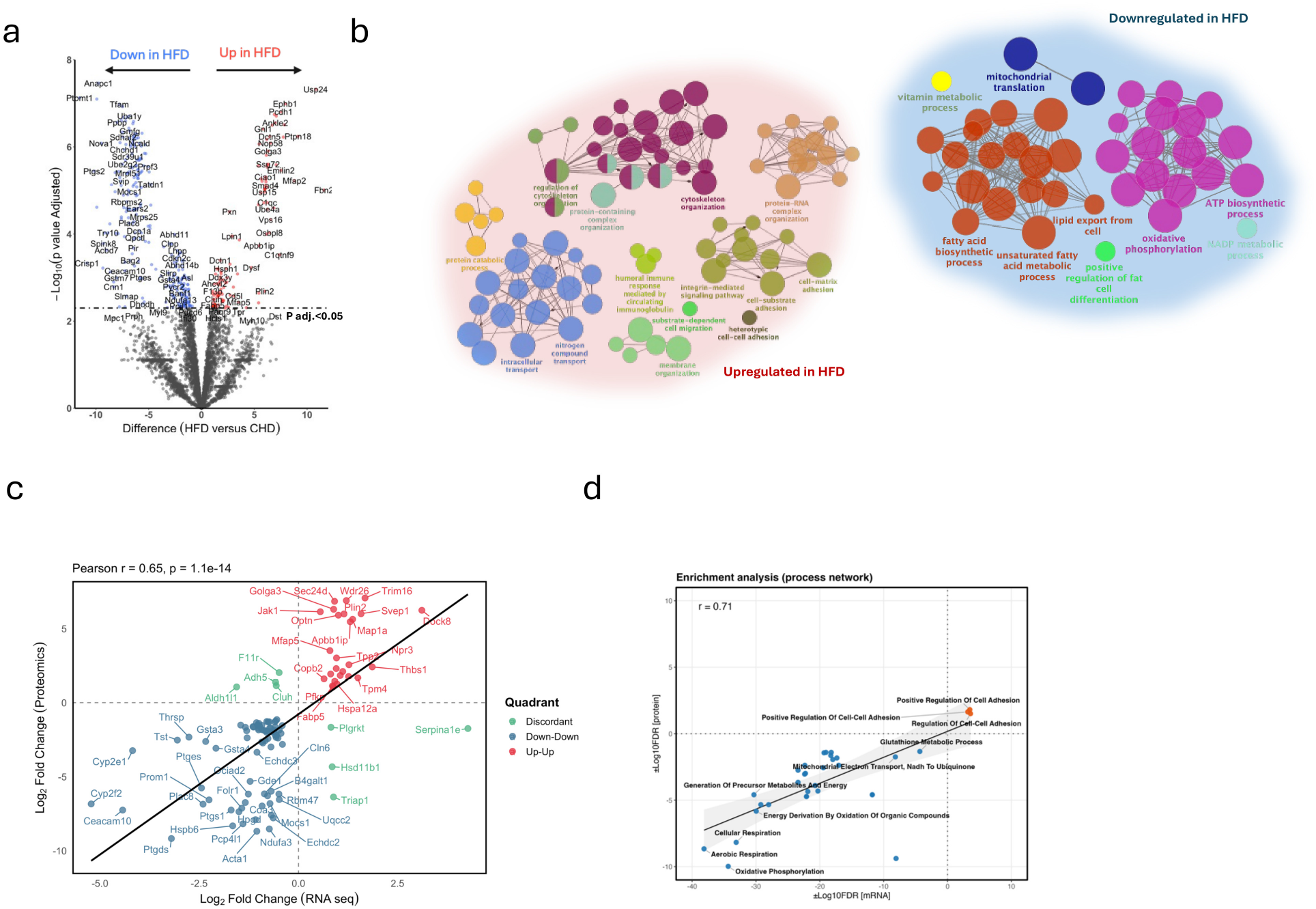
Proteomic and integrated multi-omics characterization of HFD-induced remodeling in visceral adipose tissue. (a) Volcano plot of all DAPs in visceral adipose tissue from HFD versus CHD mice, showing log₂ fold change versus adjusted p value and highlighting significantly upregulated (red) and downregulated (blue) proteins. (b) Network visualization of enriched biological processes among upregulated (bottom, colored clusters) and downregulated (top, blue/gray clusters) genes in HFD, showing functionally related process modules. (c) Correlation plot comparing RNA-seq and proteomics log₂ fold changes (HFD versus CHD) across matched gene-protein pairs, with concordantly upregulated pairs highlighted in red, concordantly downregulated pairs in blue, and discordant RNA-protein pairs in gray. (d) Enrichment analysis of concordantly regulated RNA-protein pairs, highlighting biological processes coordinately regulated at both the transcript and protein levels in HFD visceral adipose tissue.

**Figure S3.**
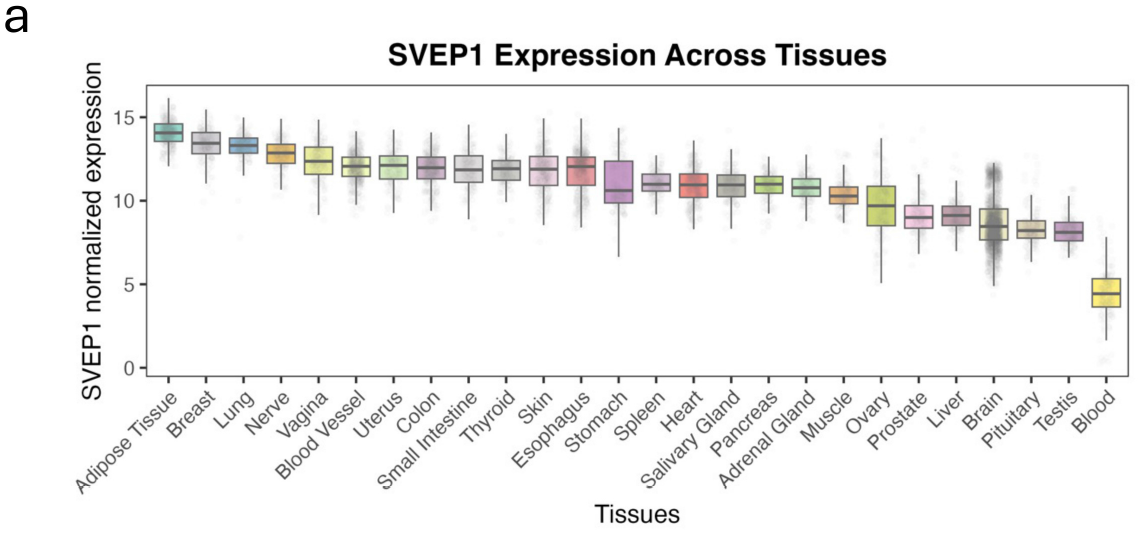
Cross-tissue characterization of *SVEP1* expression patterns. (a) *SVEP1* mRNA expression across more than 30 human tissues in the GTEx dataset, shown as normalized expression values per tissue. Data are descriptive (per-tissue GTEx medians).

**Figure S4.**
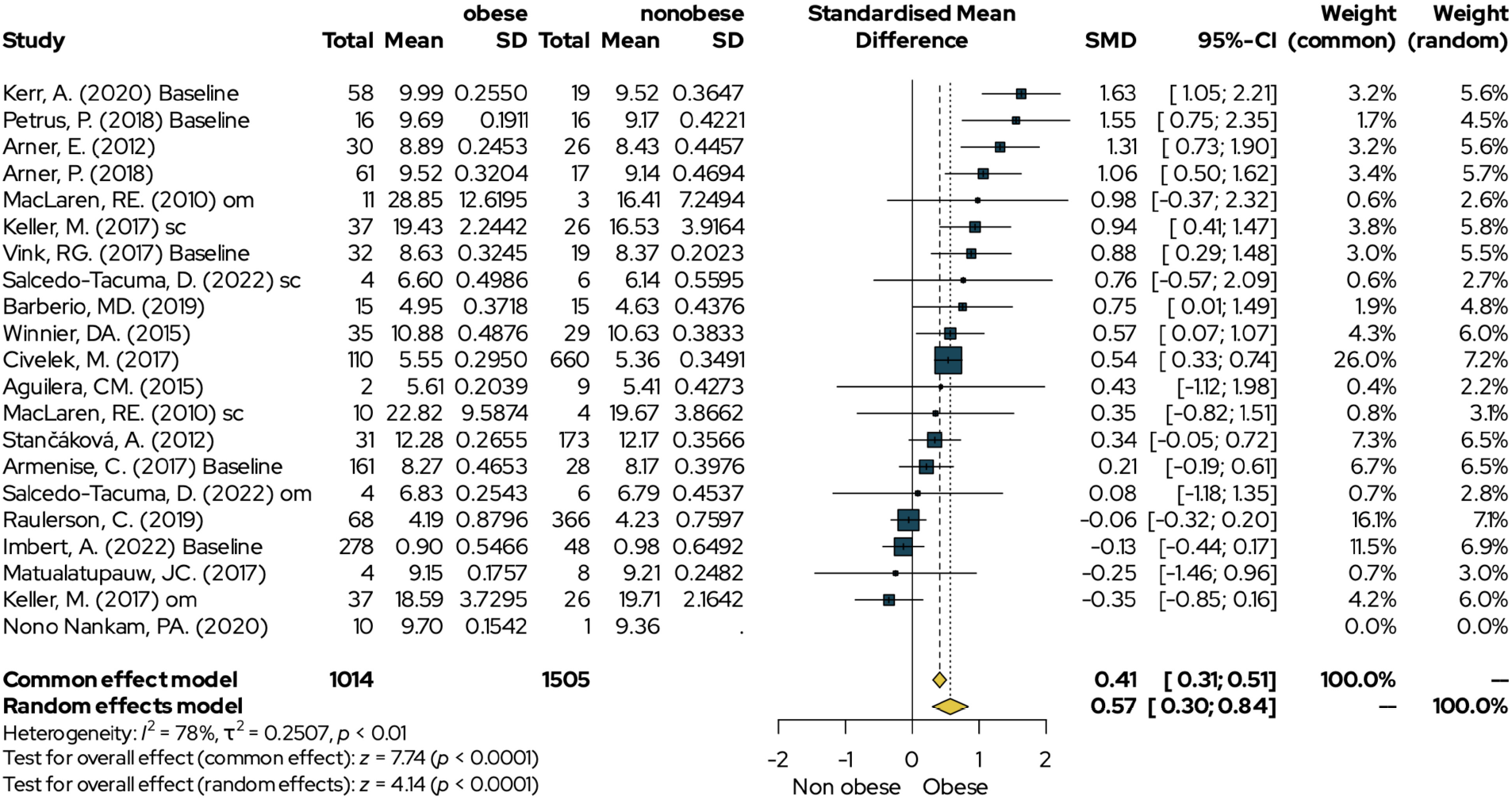
Meta-analysis of *SVEP1* expression in obese versus non-obese human adipose tissue. Forest plot displaying SMD of *SVEP1* expression between obese (n = 1,014 total) and non-obese (n = 1,505 total) subjects across 20 independent studies. Individual study effect sizes with 95% confidence intervals (CI) are shown as squares with horizontal lines. Diamond at the bottom represents the pooled effect estimate under common-effect (SMD = 0.41 [0.31; 0.51]) and random-effects (SMD = 0.57 [0.30; 0.84]) models. Heterogeneity: I² = 78%, τ² = 0.2507, p < 0.01. Test for overall effect (common effect): z = 7.74 (p < 0.0001); test for overall effect (random effects): z = 4.14 (p < 0.0001). Weights for each model are indicated.

**Figure S5.**
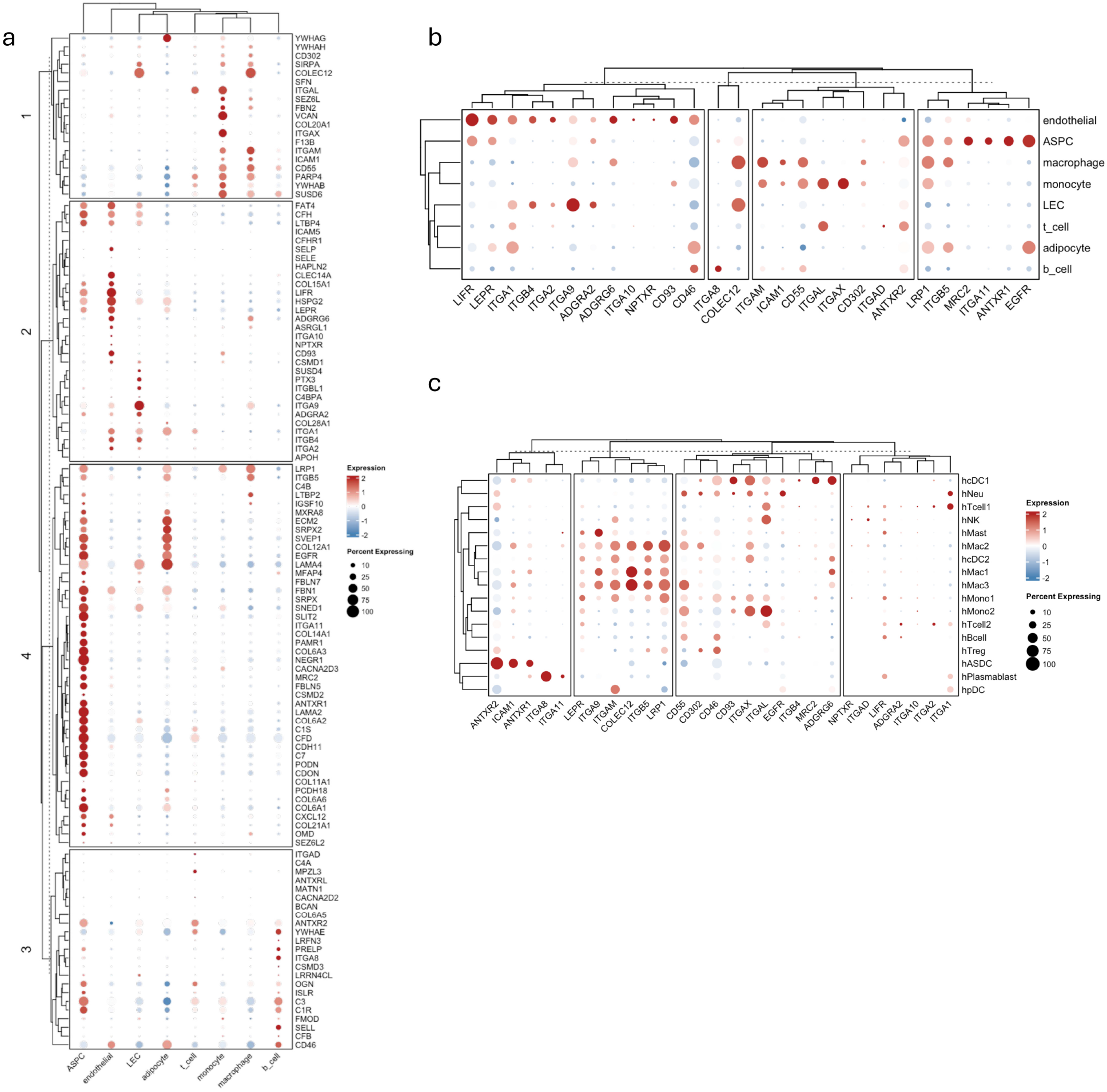
Cell-type-resolved expression patterns of the SVEP1 interactome and receptor landscape in human adipose tissue. (a) Dot-plot visualization of expression patterns of the full SVEP1 interactome across human adipose tissue cell types using Emont et al. adipose single-cell RNA-seq, organized into four major expression clusters. Cell types include endothelial cells, ASPCs, macrophages, monocytes, LECs, T cells, adipocytes, and B cells. Dot size represents the percentage of expressing cells, and color scale reflects normalized expression. (b) Dot-plot showing expression patterns of the SVEP1-associated receptor set across human adipose tissue cell types, clustered by gene and cell type to reveal coordinated receptor modules. (c) Subset dot plot showing expression patterns of SVEP1-associated receptors across immune cell populations (hcDC1, hNeu, hTcell1, hNK, hMast, hMac2, hcDC2, hMac1, hMac3, hMono1, hMono2, hTcell2, hBcell, hTreg, hASDC, hPlasmablast, hpDC). Dot size represents the percentage of expressing cells, and color reflects normalized mean expression.

**Figure S6.**
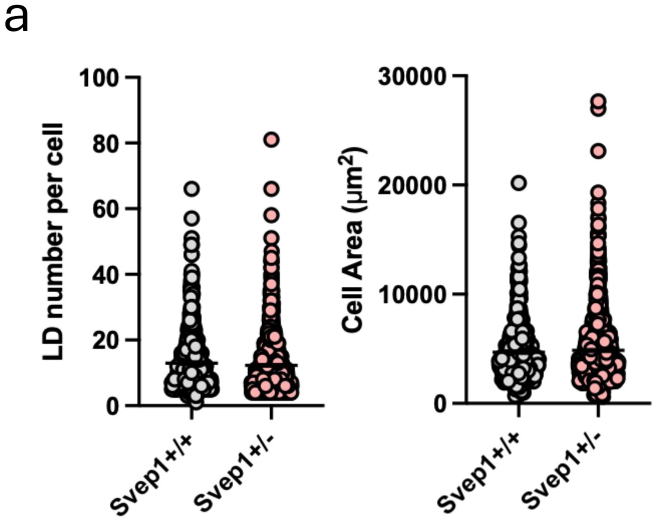
Influence of SVEP1 on adipocyte differentiation and lipid droplet morphology in vitro. (a) Quantification of lipid droplet number per field and cell area. A total of 11–14 fields per condition were analyzed from three independent biological replicates. Statistical significance was calculated using one-way ANOVA, data are presented as mean ± SEM.

**Figure S7.**
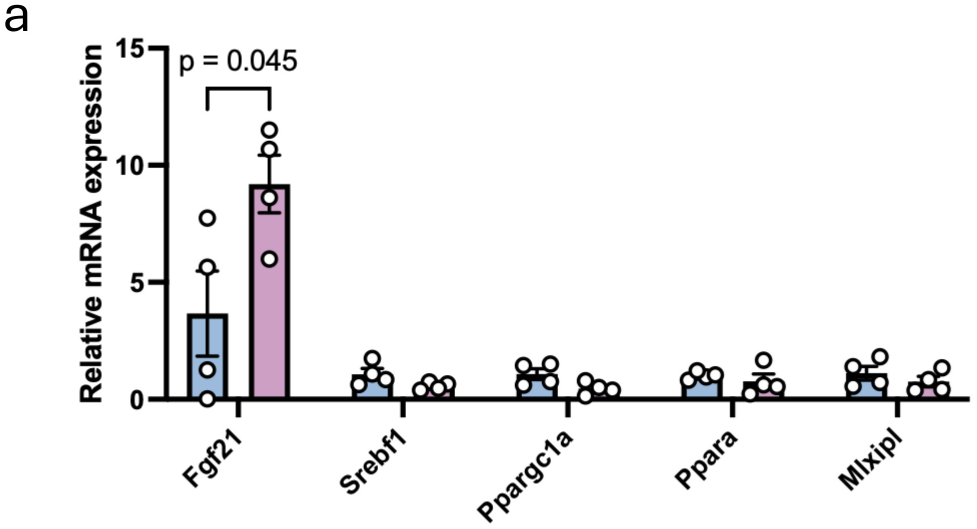
qPCR of liver genes. (a) qPCR analysis of liver gene expression from HFD-fed *Svep1*+/+ and *Svep1*+/– mice (n = 4 per group). Statistical significance was calculated using two-tailed unpaired Student’s t-test; data are presented as mean ± SEM.

**Figure S8.**
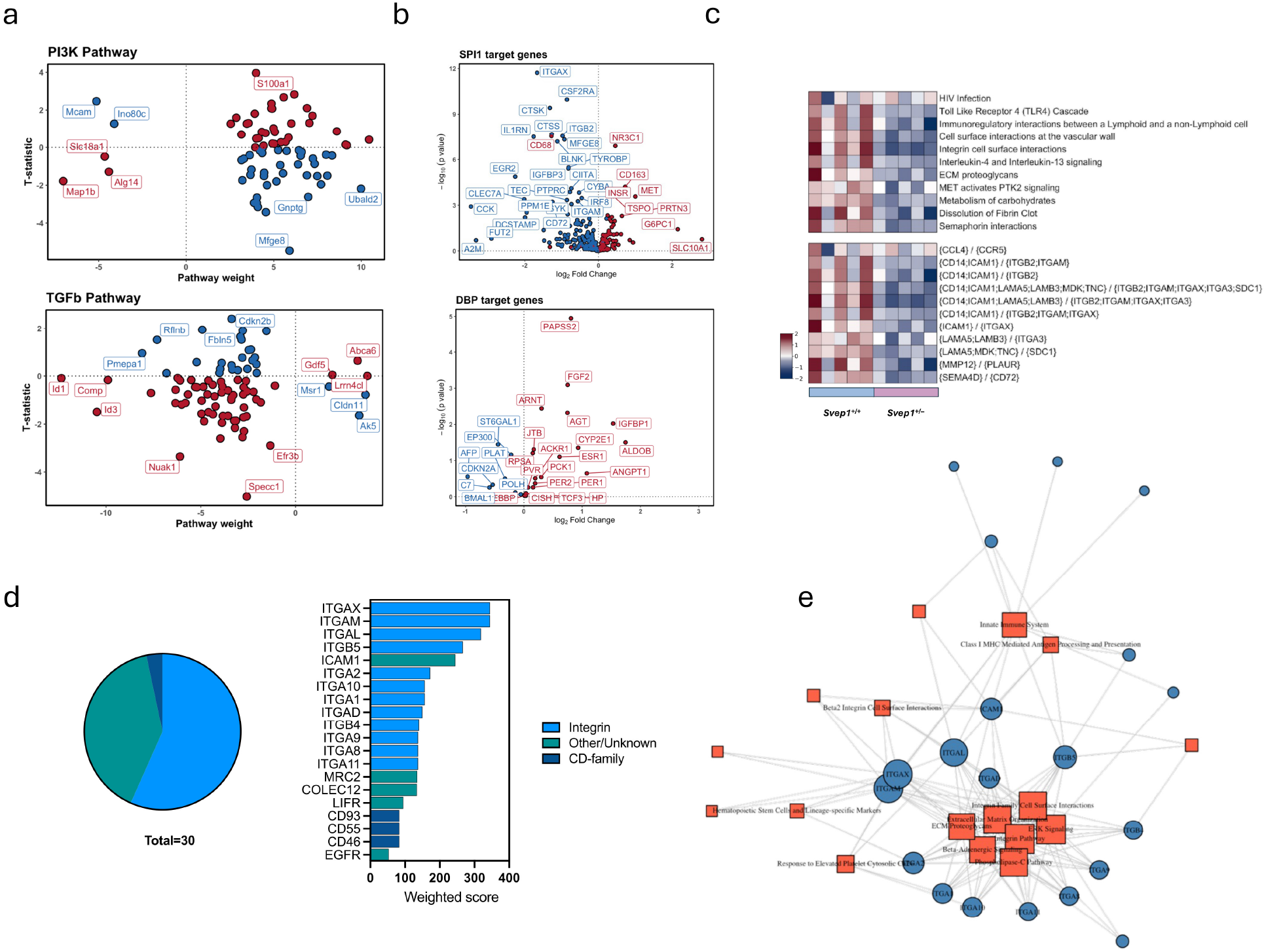
Integrated pathway, transcription factor, ligand-receptor, and ECM-oriented analyses of visceral adipose tissue from HFD-fed. *Svep1*+/+ **and** *Svep1*+/– **mice.** (a) PROGENy pathway analysis of epididymal adipose tissue from HFD-fed *Svep1*+/+ and *Svep1*+/– mice, highlighting genes with the strongest positive (red) or negative (blue) contributions to the top upregulated and top downregulated signaling pathways (PI3K pathway, top; TGF-β pathway, bottom) in the *Svep1*+/– state. (b) Volcano plots of transcription factor target genes corresponding to the most differentially active transcription factors identified by decoupleR/DoRothEA (SPI1 target genes, top; DBP target genes, bottom), showing significantly regulated downstream targets in *Svep1*+/– versus *Svep1*+/+ HFD samples. (c) BulkSignalR ligand-receptor analysis comparing *Svep1*+/– and *Svep1*+/+ HFD adipose tissue, showing pathway-level regulation inferred from ligand-receptor interactions and identifying signaling programs enriched or suppressed upon reduced *Svep1*. The heatmap summarizes receptor-pathway associations with ligand-receptor complexes listed. (d) Weighted receptor family-and receptor-level involvement analysis integrating DEG-derived pathways with the curated SVEP1 receptor set. The pie chart shows receptor-family distribution, and the bar plot shows individual receptor weighted scores colored by family. (e) Network representation of the integrated SVEP1 receptor-pathway landscape, showing receptor nodes, pathway nodes, and their functional connections.

**Figure S9.**
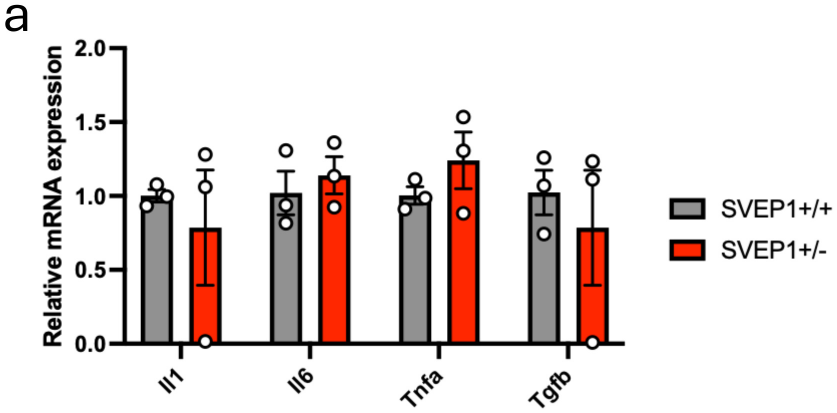
Inflammatory cytokine expression in *Svep1+/– adipocytes following* LPS stimulation. (a) qPCR analysis of proinflammatory genes in *Svep1*+/+ (gray) and *Svep1*+/– (red) adipocytes following 8 h of LPS stimulation (n = 3). Statistical significance was calculated using a two-tailed unpaired Student’s t-test; data are presented as mean ± SEM.

## Supplementary Tables

**Table S1.**
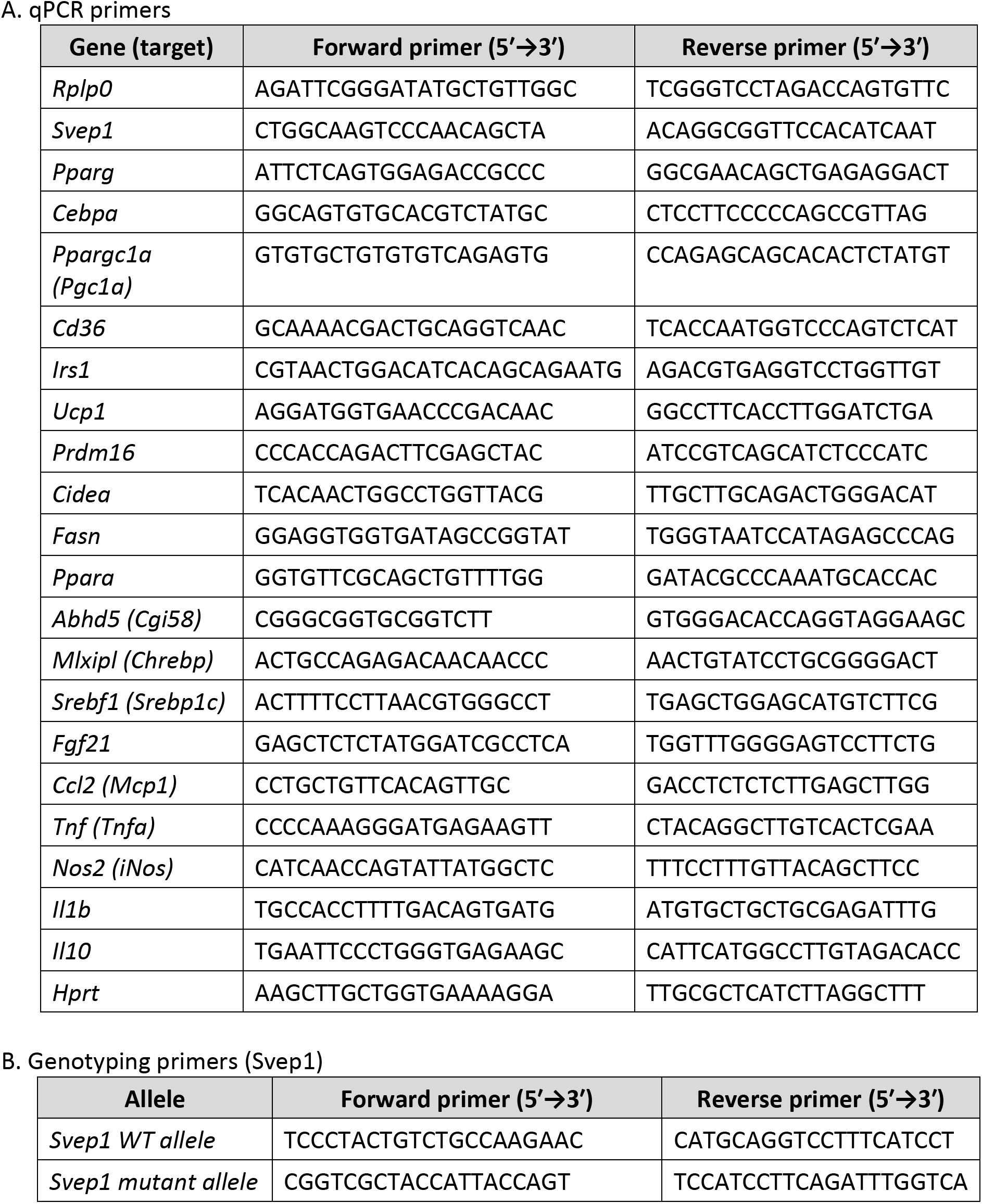
Oligonucleotide primer sequences used in this study.

**Table S2.** Antibodies used for flow cytometry.

| Target | Fluorophore | Clone | Catalog no. | Isotype | Vol. / test | Vendor |
| --- | --- | --- | --- | --- | --- | --- |
| *CD45* | AF700 | 30-F11 | 103128 | Rat IgG2b, $\kappa$ | 0.8 μl | BioLegend |
| *CD45* | APC | 30-F11 | 07512-80 | Rat IgG2b | 1 μl | Biogems /<br>PeproTech |
| *F4/80* | FITC | BM8.1 | 02922-80 | Rat IgG2a | 1 μl | Biogems /<br>PeproTech |
| *CD11b* | PE/Cy5 | M1/70 | 03221-65 | Rat IgG2a | 1 μl | Biogems /<br>PeproTech |
| *MHCII* | PE/Cy7 | M5/114.15.2 | 107629 | Rat IgG2b, $\kappa$ | 0.8 μl | BioLegend |
| *CD11c* | APC/Cy7 | N418 | 117323 | Armenian<br>hamster IgG | 0.8 μl | BioLegend |
| *ITGA9* | PE | Tyr31-Val979 | FAB3827P | Goat IgG | 10 μl | R&D Systems |

